# Aurora B and Condensin are dispensable for chromosome arm and telomere separation during meiosis II

**DOI:** 10.1101/2020.01.07.897033

**Authors:** Julien Berthezene, Céline Reyes, Tong Li, Stéphane Coulon, Pascal Bernard, Yannick Gachet, Sylvie Tournier

## Abstract

In mitosis, while the importance of kinetochore-microtubule attachment has been known for many years, increasing evidence suggests that telomere dysfunctions also perturb chromosome segregation by contributing to the formation of chromatin bridges at anaphase. Recent evidence suggests that Aurora B ensures proper chromosome segregation during mitosis not only by controlling kinetochore-microtubule attachment but also by regulating telomere and chromosome arm separation. However, whether and how Aurora-B governs telomere separation during meiosis has remained unknown. Here, we show that fission yeast Aurora B localizes at telomeres during meiosis I and promotes telomere separation independently of the meiotic cohesin Rec8. In meiosis II, Aurora-B controls kinetochore-microtubule attachment but appears dispensable for telomere and chromosome arm separation. Likewise, condensin activity is nonessential in meiosis II for telomere and chromosome arm separation. Thus, in meiosis, the requirements for Aurora-B are distinct at centromeres and telomeres, illustrating the critical differences in the control of chromosome segregation between mitosis and meiosis II.

## INTRODUCTION

In eukaryotes, accurate chromosome segregation during mitosis relies on the migration of sister-chromatids towards the opposite poles of the mitotic spindle. To achieve correct chromosome segregation, kinetochores (KTs), multiprotein complexes assembled at centromeres, must interact with microtubules (MTs). When sister KTs are captured by MTs coming from opposite poles of the mitotic spindle, the mitotic chromosome becomes bilaterally attached and congresses towards the spindle equator. At anaphase sister chromatids separate and move towards the poles, due to the proteolytic cleavage of cohesin, which ensures sister chromatid cohesion (Uhlmann et al., 1999). Sister chromatid cohesion defines the correct back-to-back arrangement of sister KTs and prevents chromosome attachment defects such as merotelic attachment, in which one KT is attached to both poles (Courtheoux et al., 2009; Gregan et al., 2011; Gregan et al., 2007; Sakuno et al., 2009). The stiffness of chromosome arms, conferred by condensin, transmits the traction forces exerted on centromeres by MTs throughout chromosome arms up to the telomeres, allowing the full separation of chromosomes (Renshaw et al., 2010). Spindle lengthening (anaphase B) further separates the two sets of chromosomes and cytokinesis ensures a complete partitioning of the two daughter cells.

It is crucial for the accurate segregation of chromosomes that cohesion is not destroyed until all chromosomes are bilaterally attached to the mitotic spindle. This key coordination is ensured by an error detection and correction system, which involves Aurora B kinase (Chan and Botstein, 1993; Lampson and Cheeseman, 2011) and the spindle assembly checkpoint (SAC) (Cleveland et al., 2003; Rieder et al., 1995; Rudner and Murray, 1996). Aurora B and SAC components operate at the level of KTs and monitor chromosome attachment, allowing the destabilization of erroneous attachments. Aurora B destabilizes faulty KT-MT attachments through the phosphorylation of key KT substrates (Cimini et al., 2006; Kelly and Funabiki, 2009; Tanaka et al., 2002). In anaphase, correction of attachment defects by Aurora B is dependent on forces generated at the spindle mid-zone during spindle elongation (Courtheoux et al., 2009; Gay et al., 2012). In addition, Aurora B regulates key mitotic and meiotic events such as, condensin binding and chromosome condensation, chromosome orientation, central spindle formation and the regulation of furrow ingression and abscission (Kotwaliwale et al., 2007; Meyer et al., 2013; Mora-Bermudez et al., 2007; Sampath et al., 2004; Steigemann et al., 2009; Tada et al., 2011). Aurora B is the catalytic subunit of the chromosomal passenger complex (CPC). In *S. pombe*, components of the CPC, Bir1/Survivin, Pic1/INCENP, and Ark1/Aurora B are major regulators of mitosis (Leverson et al., 2002; Morishita et al., 2001; Petersen et al., 2001). Inhibition of Aurora B using a Shokat analogue-sensitive mutant leads to chromosome segregation defects (Gay et al., 2012; Hauf et al., 2007). Several mitotic substrates of Aurora B have been identified (Carmena et al., 2012; Koch et al., 2011). Notably, Aurora B regulates the binding of condensin to chromosomes in a wide range of systems (Giet and Glover, 2001; Morishita et al., 2001; Ono et al., 2004). In fission yeast, Aurora B-dependent phosphorylation of Cnd2, the kleisin subunit of condensin, promotes condensin recruitment to chromosomes (Nakazawa et al., 2011; Tada et al., 2011). Similarly, in human cells, phosphorylation of the kleisin CAP-H by Aurora B promotes efficient association of condensin I to mitotic chromosomes (Lipp et al., 2007; Ono et al., 2004; Tada et al., 2011).

In line with its wide range of functions, Aurora B exhibits a dynamic localization during the cell cycle. Aurora B concentrates at the inner centromeric chromatin in metaphase, and eventually relocalizes at the spindle midzone during anaphase (Petersen et al., 2001). Aurora B has also been observed at telomeres during mitosis (Reyes et al., 2015; Vanoosthuyse et al., 2007; Yamagishi et al., 2010). Recent studies described a key role for Aurora B as a regulator of telomere structural integrity. In mammalian cells, loss of Aurora B function affects TERF1/TRF1 (a component of the telomeric Shelterin complex) resulting in aberrant telomere structure (Chan et al., 2017). Aurora B also regulates an active telomere conformational change from looped to linear during mitotic arrest, coincident with ATM-dependent telomere DNA damage response activation (Van Ly et al., 2018). Moreover, it has been proposed that mammalian telomeres exist in an under-protected state during mitosis due to an Aurora B–dependent effect on the telomeric Shelterin complex (Hayashi et al., 2012; Orthwein et al., 2014).

We previously identified a Telomere Disjoining Pathway (or TDP) that governs the efficient separation of chromosome ends during mitosis, and which involves an Aurora-B dependent cascade of events at fission yeast telomeres (Reyes et al., 2015). In mitosis, the physical separation of telomeres is a tightly regulated, two step process, which relies on telomeric proteins, condensin and Aurora B kinase. Aurora B primes telomeres for separation in metaphase, by delocalizing several proteins including Shelterin, Swi6^HP1^ and cohesin, and it promotes telomere separation in anaphase by facilitating condensin loading (Reyes et al., 2015).

The emerging roles of Aurora B in mitosis at those chromosomal loci, ie. telomeres and chromosome arms, prompted us to investigate the function of this kinase during the meiotic process. Indeed, meiosis represents an attractive model in which to study the intricate relationship between telomeres, condensin and Aurora B since it involves a single round of DNA replication followed by two series of chromosome segregation. Meiosis I relies on the pairing of replicated homologous chromosomes and on the pulling of pairs of homologous sister centromeres in opposite directions while in meiosis II, sister centromeres split as they do in mitosis. In addition, during meiosis, telomeres exhibit critical and striking features. In pre-meiotic S-phase, chromosomes reorganize to form the bouquet configuration where telomeres are arranged as a large cluster beneath the spindle pole body (Chikashige et al., 1994; Scherthan, 2001). This telomere cluster is tightly anchored to the nuclear envelope (Chikashige et al., 2006; Hiraoka and Dernburg, 2009). Previous studies in *S. pombe* have established that the telomere bouquet has crucial functions in the preparation of meiotic chromosome segregation. These functions encompass promoting homologous chromosome recombination (Cooper et al., 1998), setting up spindle formation (Tomita and Cooper, 2007), participating in meiotic centromere assembly (Klutstein et al., 2015) and finally controlling prophase progression and exit (Moiseeva et al., 2017).

In the present study, we used live single cell imaging to analyze centromere, telomere and chromosome arm separation when Aurora B or condensin were selectively inactivated in meiosis I or meiosis II. We demonstrate that the activity of Aurora B and condensin are required for telomere and chromosome arm separation in meiosis I but not in meiosis II. We further show that the functions of Aurora B in KT-MT attachment and telomere separation can be distinguished. Finally, our study also suggests the existence of a non-canonical mechanism to segregate chromosome arms in meiosis II when Aurora or condensin are inactivated.

## RESULTS

### Telomere separation is a two steps process in meiosis I as opposed to meiosis II

To investigate the mechanisms controlling telomere separation in meiosis, we performed live imaging of cells expressing a spindle pole body (SPB) marker, Cdc11-CFP (Tournier et al., 2004) and a telomere marker protein, Taz1-GFP; a homologue of mammalian TRF1/TRF2 that binds exclusively to telomere repeats (Chikashige and Hiraoka, 2001; Cooper et al., 1997; Vassetzky et al., 1999) (Fig. 1).

**Figure 1:**
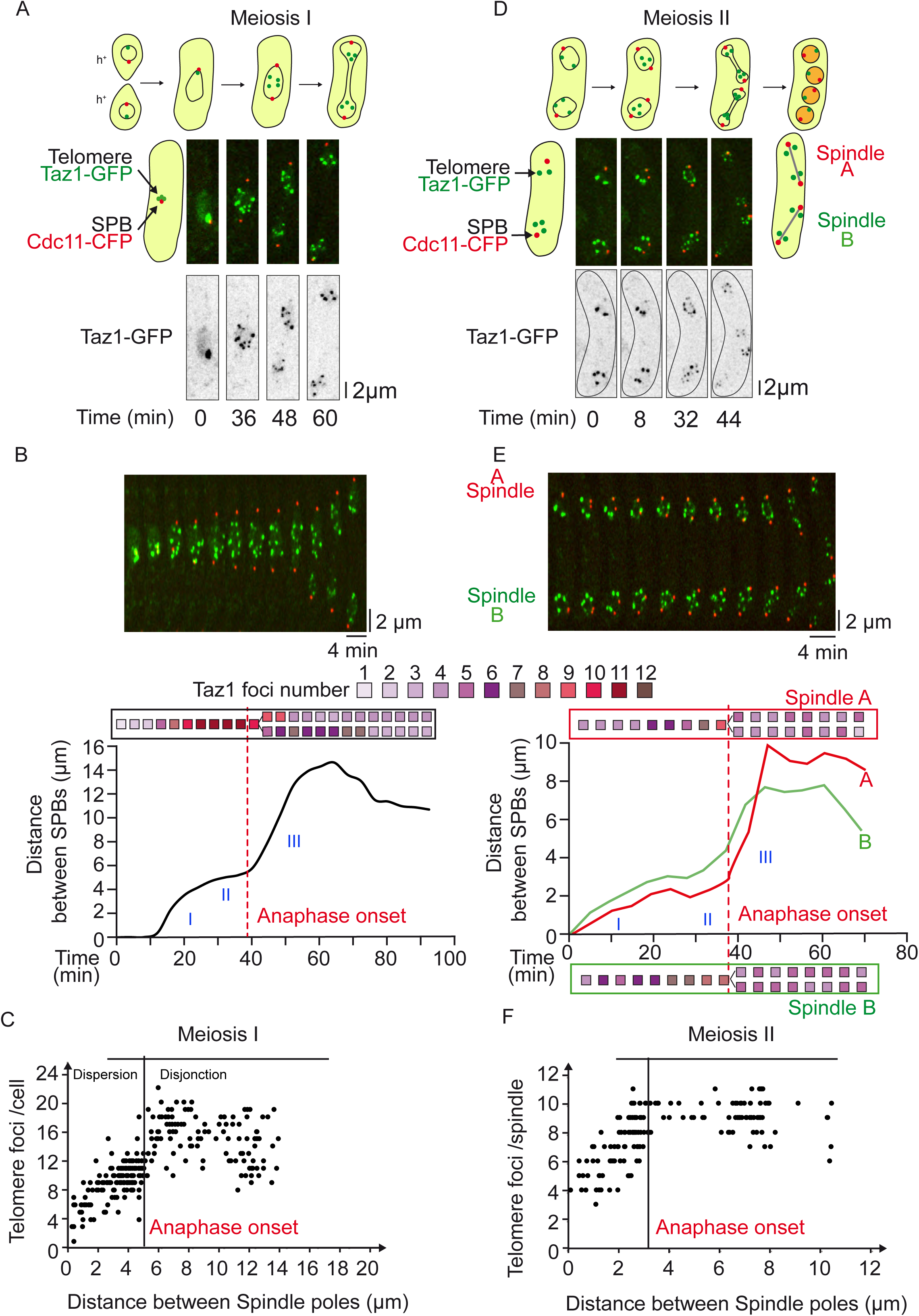
Telomere cluster dissociation occurs in two steps in meiosis I but in a single step during metaphase of meiosis II. **A**. Selected frames from a movie of a live cell undergoing meiosis I. Telomeres are visualized via Taz1-GFP (green) and SPBs via Cdc11-CFP (red). **B**. Telomere foci dynamics during meiosis I. A kymograph and analysis of the movie shown in A are presented. The number of telomere foci is shown with a color code for each frame of the movie. After anaphase I onset, telomere foci numbers were counted separately in both sets of segregated chromosomes. **C**. Number of telomere foci according to the distance between SPB foci. Data were measured on static images of cells (n = 264) in meiosis I. Characteristic spindle length of anaphase I onset was estimated to be in average of 5µm. **D**. Selected frames from a movie of a live cell undergoing meiosis II. Telomeres are visualized via Taz1-GFP (green) and SPBs via Cdc11-CFP (red). **E**. Telomere foci dynamics during meiosis II. A kymograph and analysis of the movie shown in D is presented, the upper and lower meiosis II spindles of the movie are defined as spindle A and spindle B respectively. The number of telomere foci is shown with a color code for each frame of the movie for both Spindle A (red box) and Spindle B (green box). After anaphase II onset, telomere foci numbers were counted separately for both sets of segregated chromosomes. **F**. Number of telomere foci according to the distance between SPB foci. Data were measured from meiosis II movies (n = 12). Characteristic spindle length of anaphase II onset was estimated at 3.2 µm.

Fission yeast has three chromosomes for 12 telomeres in mitosis. At the beginning of meiosis, during the horse-tail movement, the twenty four telomeres are organized in clusters at the SPBs while the centromeres are positioned away from the SPBs (bouquet conformation). We followed individual *taz1-gfp cdc11-cfp* cells through meiosis I (Fig.1A-C). In early meiosis I, a short spindle (up to 3.0 μm) is formed, as judged by SPB separation. A second phase (from 3 to 5.0 μm) is observed where spindle length is roughly maintained until full separation of homologous chromosomes (anaphase A). The third phase of meiosis I consists entirely of anaphase B, during which the spindle elongates along the long axis of the cell (Fig.S1C, movie 1). Telomeres separate in two steps during meiosis I. Firstly, telomere cluster dispersion is initiated in early meiosis I prior to anaphase A (from 1 to 12 foci; spindle size from 0 to 5 μm), and secondly, clusters fully dissociate during anaphase B when chromosome arms separate (telomere disjunction, from 10 to 20 foci, spindle size from 5 to 15 μm, movie 1). An example of meiosis I telomere dispersion and disjunction is shown in a single cell analysis (Fig. 1B) together with a quantification of Taz1 foci during meiotic progression in multiple individual meiotic cells (Fig. 1C) confirming that telomere separation is a two step process. Importantly, during prophase of meiosis II, telomeres were able to form new clusters (Fig. S1A). Quantification of Taz1 focus number and position confirmed that the re-establishment of telomere clusters and their re-localization to the nuclear periphery occurs after spindle disassembly in prophase II before the onset of metaphase II (Fig. S1B).

We similarly analyzed telomere separation during meiosis II (Fig.1 D-F). In early meiosis II, two short spindles are formed (up to 2.0 μm), as judged by SPB separation. A second phase (from 2 to 3.0 μm) is observed where both spindles roughly maintained the same size until sister chromatids separate (anaphase A). The third phase consists of anaphase B, during which the two spindles moderately elongate within the cell (up to 10 μm) (Fig.S1 D, movie 2). An example of meiosis II telomere dissociation is shown in a single cell analysis (Fig. 1E) together with a quantification of Taz1 foci during meiosis II progression in several cells (Fig.1F). Interestingly, we observed a single step in telomere separation, which was mostly achieved in metaphase II, *ie.* prior the full separation of sister chromatids in anaphase II (Fig. 1E and F). Quantification of Taz1 foci confirmed the re-establishment of telomere clusters during spore formation (Fig. S1A).

Together, these experiments demonstrate that telomere clustering is highly regulated throughout the meiotic cell cycle and while telomere dissociation is a two-step process in meiosis I, it occurs in a single phase in meiosis II.

### Aurora B kinase controls telomere and chromosome arm separation in meiosis I but not in meiosis II

To test whether Aurora B plays a role in telomere separation in meiosis, we used ATP analogue sensitive alleles of Aurora B (Shokat mutant *ark1-as3*) (Hauf et al., 2007). Aurora B activity was specifically inhibited in meiosis I or meiosis II (see Material and Methods) by adding the non-hydrolizable ATP analogue 1-NA-PP1 (10μM). Meiotic progression of *ark1-as3* cells expressing telomere *(taz1-gfp,* green) and SPB markers (*cdc11-cfp,* red) was recorded ten minutes after adding the inhibitor (Fig. 2). In meiosis I, single cell analysis of the number of Taz1 foci with respect to spindle size reveals that Aurora B inhibition significantly perturbed telomere separation, even after full spindle elongation (Fig.2A-C) and even promoted re-clustering of telomeres in anaphase I (Fig.2A-C; movie 3). Thus, in addition to its previously described role at centromeres in the correction of faulty KT-MT attachment (Fig.S2B-C), Aurora B is required in meiosis I for telomere separation.

**Figure 2:**
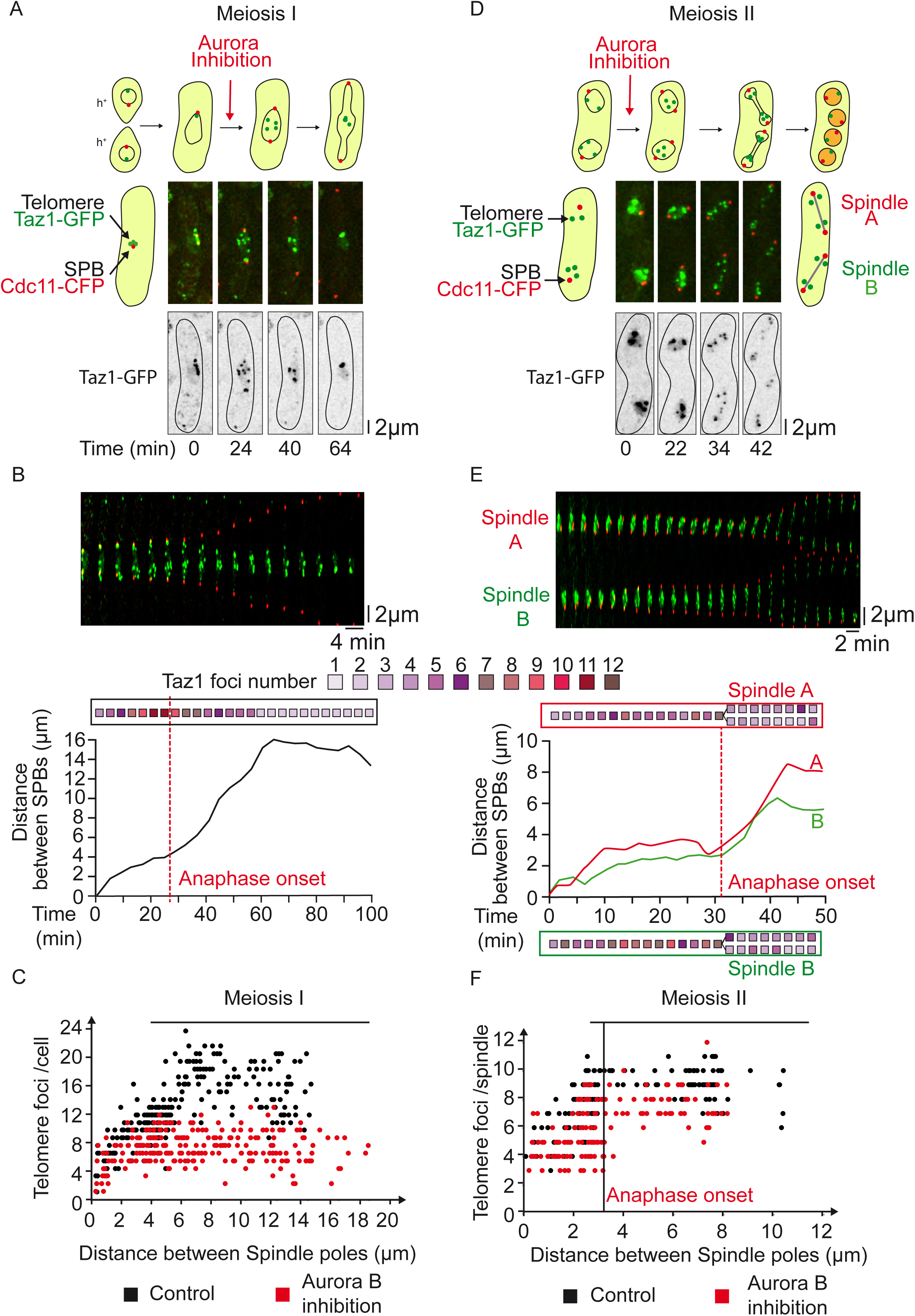
Aurora kinase function is required for the separation of telomeres in meiosis I but is dispensable in meiosis II. **A**. Selected frames from a live *ark1-as3* cell undergoing meiosis I under Aurora inhibition. Aurora inhibition is performed by exposure of an *ark1-as3* cell to 10µM 1-NA-PP1. Telomeres are visualized via Taz1-GFP (green) and SPBs via Cdc11-CFP (red). **B**. Telomere foci dynamics during meiosis I under Aurora inhibition. A kymograph and analysis of the movie shown in A is presented. The number of telomere foci is shown with a color code for each frame of the movie. After anaphase I onset, telomeres do not separate and instead gradually reform clusters. **C**. Number of telomere foci according to the distance between SPB foci. Data were measured from static images of *ark1-as3* cells (n=240) under Aurora inhibition (red dots). Data from Figure 1C were reported as a control (black dots). **D**. Selected frames from a movie of a live *ark1-as3* cell under meiosis II-specific Aurora inhibition. Telomeres are visualized via Taz1-GFP (green) and SPBs are visualized via Cdc11-CFP (red). Aurora inhibition was performed by adding 10µM 1-NA-PP1, 10min prior the start of image acquisition. **E**. Telomere foci dynamics under meiosis II-specific Aurora inhibition. A kymograph and analysis of the movie shown in D is presented, the upper and lower meiosis II spindles of the movie are defined as spindle A and spindle B respectively. The number of telomere foci is shown with a color code for each frame of the movie for both Spindle A (red box) and Spindle B (green box). After anaphase II onset, telomere foci number was counted separately in both segregated sets of chromosomes. **F**. Number of telomeric foci under Aurora inhibition according to the distance between SPB foci. Data were measured on meiosis II movies of *ark1-as3* cells (n = 10) under meiosis II-specific Aurora inhibition (red dots). Aurora inhibition was performed by adding 10µM 1-NA-PP1, 10min prior the start of image acquisition. Data from Figure 1F was also reported as a control (black dots).

In this set of experiments, we noticed that when Aurora inhibition was conducted from the beginning of meiosis I, the onset of meiosis II was also deeply perturbed because of spindle collapse occurring in meiosis I (not shown). Thus, to specifically analyze the role of Aurora B during meiosis II, we added the inhibitor 1-NA-PP1 to *ark1-as3 taz1-gfp cdc11-cfp* cells only during prophase II and followed cells progressing into meiosis II (see Material and Methods; Fig.2 D-F). An example of telomere separation during meiosis II when Aurora is inhibited is shown in a single cell analysis (Fig. 2D-E, movie 4) together with a quantification of Taz1 foci during meiosis II progression in multiple cells (Fig.2F). Surprisingly, when Aurora was inhibited specifically in meiosis II, the kinetic of telomere separation remained unchanged. Telomeres appeared clustered during metaphase II and fully separated at the anaphase II transition (Fig.2E-F). This observation was not due to the lack of efficiency of Aurora inhibition since we found that centromere segregation was greatly impacted by Aurora inhibition in meiosis II (Fig.S2D and E). Accordingly, using histone H3-mRFP (red) and Taz1-GFP (green) markers, we observed that Aurora inhibition blocked the separation of chromosome arms and telomeres in meiosis I (Fig.3A-B). In sharp contrast, although lagging chromosomes were present, chromosome arms and telomeres successfully separated during anaphase II despite the inhibition of Aurora B (Fig.3C-D).

**Figure 3:**
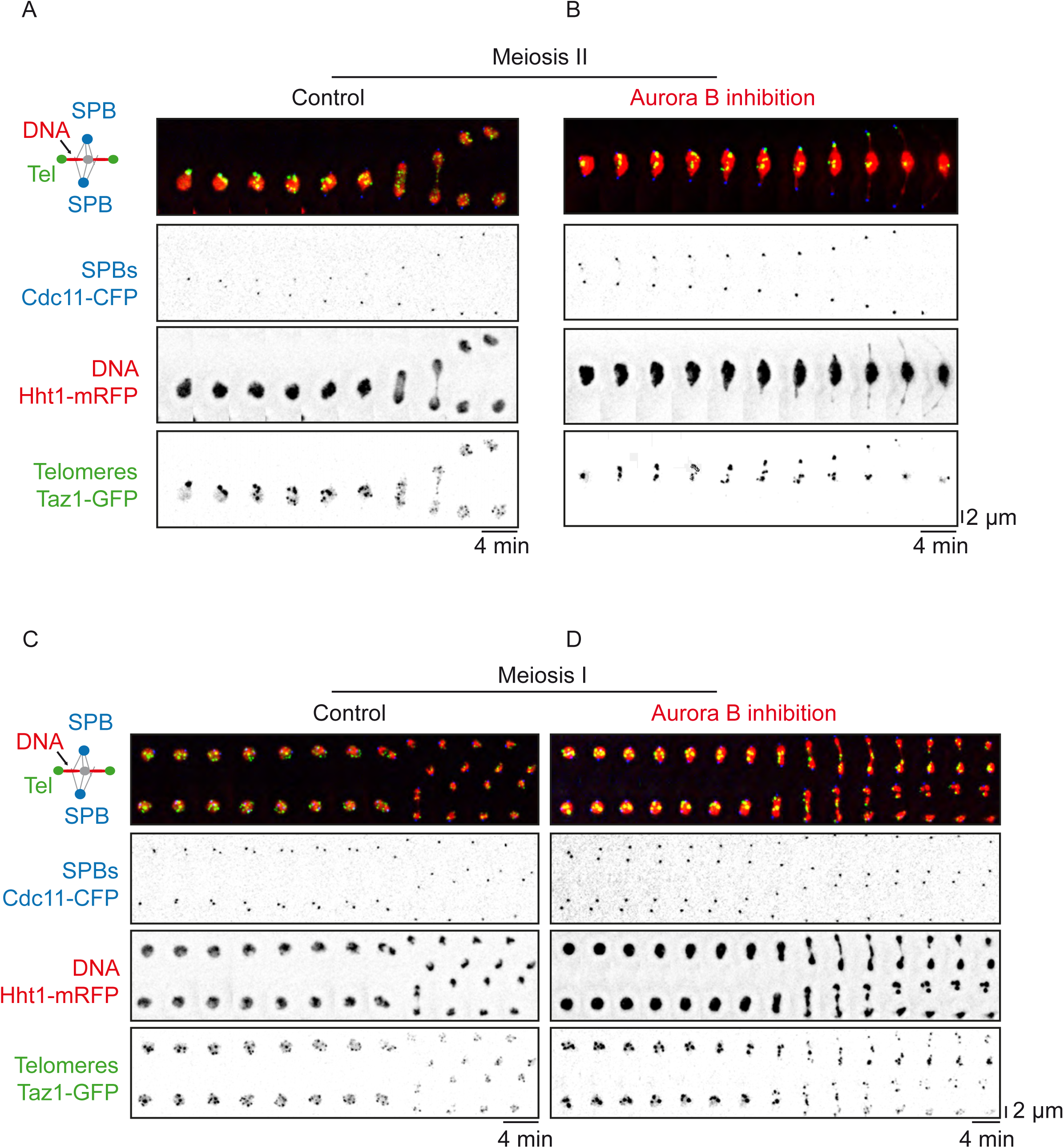
Aurora kinase is required for the separation of chromosome arms in meiosis I but not in meiosis II. Time lapse imaging of chromosome arms disjunction in live meiotic *ark1-as3* cells. SPBs are visualized via Cdc11-CFP (blue), chromosome arms via Hht1-mRFP (histone H3, red) and telomeres via Taz1-GFP (green). Frames were acquired at intervals of 4min. **A**. Example of a cell during meiosis I progression. **B**. Example of a cell during meiosis I progression in the presence of 10µM 1-NA-PP1. **C**. Example of a cell during meiosis II progression. **D**. Example of a cell during meiosis II progression in the presence of 10µM 1-NA-PP1.

To better understand the differences between Aurora’s role in equational (meiosis II) versus reductional (meiosis I) chromosome segregation, we performed imaging of live cells simultaneously expressing markers for the SPBs-Kinetochores (Cdc11-CFP, Ndc80-CFP, blue), telomeres (Taz1-RFP, red) and Aurora B (Ark1-GFP, green) (Fig.4). In addition to its well-known localization at KTs and the spindle midzone in metaphase of meiosis I and II (Hauf et al., 2007; Petersen et al., 2001), we observed that Aurora B also co-localizes transiently to telomeres in metaphase of meiosis I (Fig.4A). Intriguingly, telomere localization of Aurora B was exclusively seen in meiosis I but not meiosis II (Fig.4A). In addition, telomere localization of Aurora in meiosis I rapidly vanished at the metaphase to anaphase transition in parallel with telomere disjunction (Fig.4B; Fig. S3).

**Figure 4:**
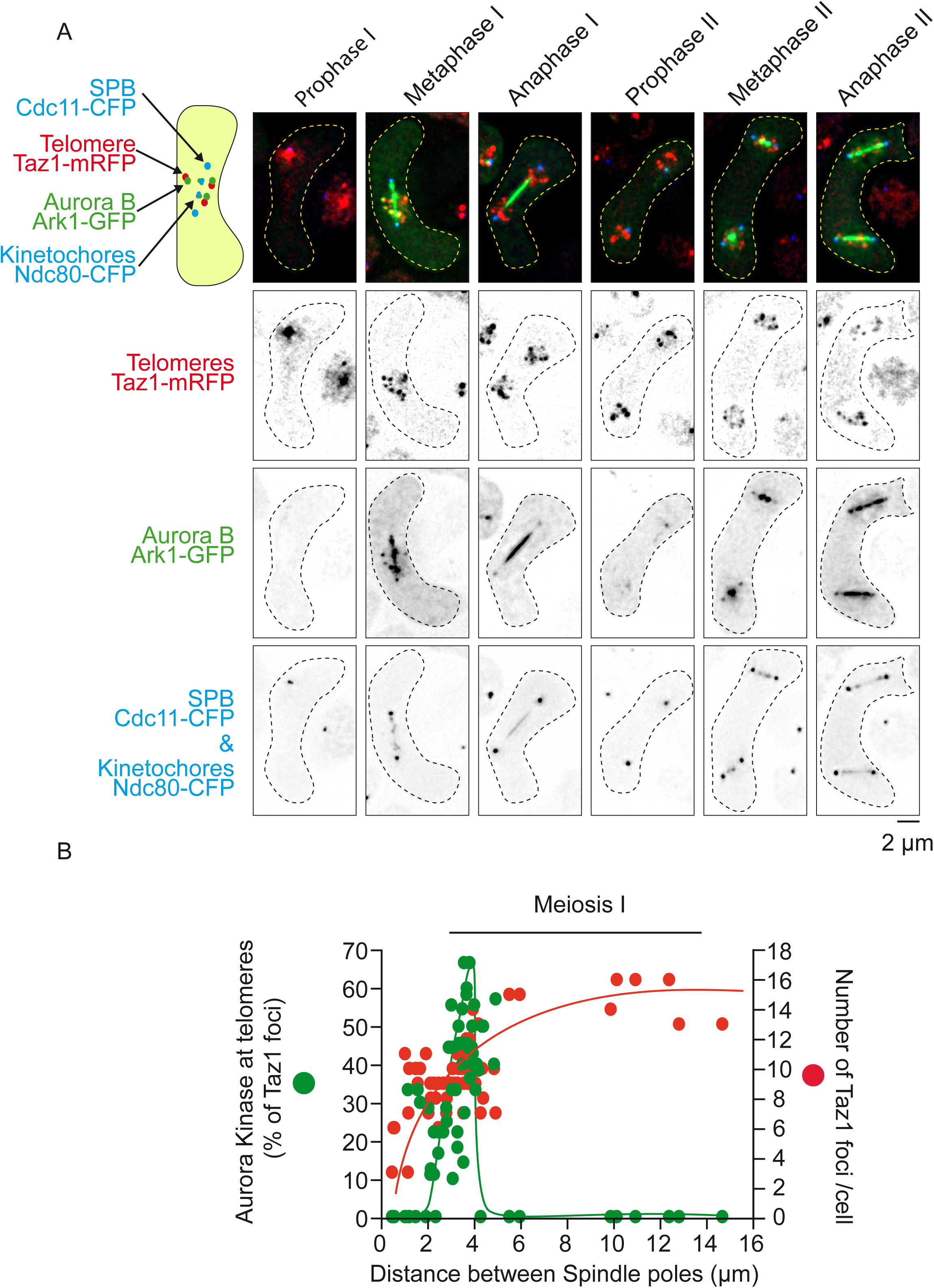
Aurora kinase localizes to telomeres during metaphase I but not during metaphase II. **A**. Static images, representative of each meiotic phase, are shown. Telomeres are visualized via Taz1-mRFP (red), Aurora via Ark1-GFP (green), SPBs via Cdc11-CFP (blue) and KTs via Ndc80-CFP (blue). **B**. Dynamics of Aurora recruitment to telomeres during meiosis I. Both telomere foci number (Taz1-mRFP, red dots) and number of Aurora foci (Ark1-GFP, green dots) localized at telomeres were quantified on static images of cells undergoing meiosis I (n = 67).

In conclusion, Aurora B activity is required for proper KT-MT attachment, and telomere and chromosome arm separation in meiosis I. Conversely, in meiosis II, Aurora B activity is dispensable for telomere and chromosome arm separation, though it remains necessary for correct KT-MT attachment. Moreover, the role of Aurora in telomere/chromosome arm separation corroborates with its localization to telomeres, as previously observed during mitosis (Reyes et al., 2015).

### Aurora controls condensin localization to chromosome arms in meiosis I and meiosis II

In fission yeast, the Aurora B-dependent phosphorylation of the kleisin Cnd2 during mitosis promotes condensin recruitment to chromosomes (Nakazawa et al., 2011; Tada et al., 2011). To determine whether this mechanism was also conserved in meiosis, we performed imaging of living cells simultaneously expressing markers for the SPBs (Cdc11-CFP, red) and the condensin subunit Cnd2 (Cnd2-GFP, green) before or after Aurora inhibition (Fig. 5). As previously described in mitosis (Nakazawa et al., 2008), we observed that condensin localizes in meiosis I to several nuclear foci compartments, possibly corresponding to nucleolar or KT foci (Fig.5A; left panel; movie 5). At anaphase onset, condensin switches to the spindle midzone and chromosome arms but foci largely disappear. A histogram analysis of foci pixel intensities of condensin localization during meiosis I progression confirms the reorganization of condensin localization during meiotic progression (Fig.5B; left panel).

**Figure 5:**
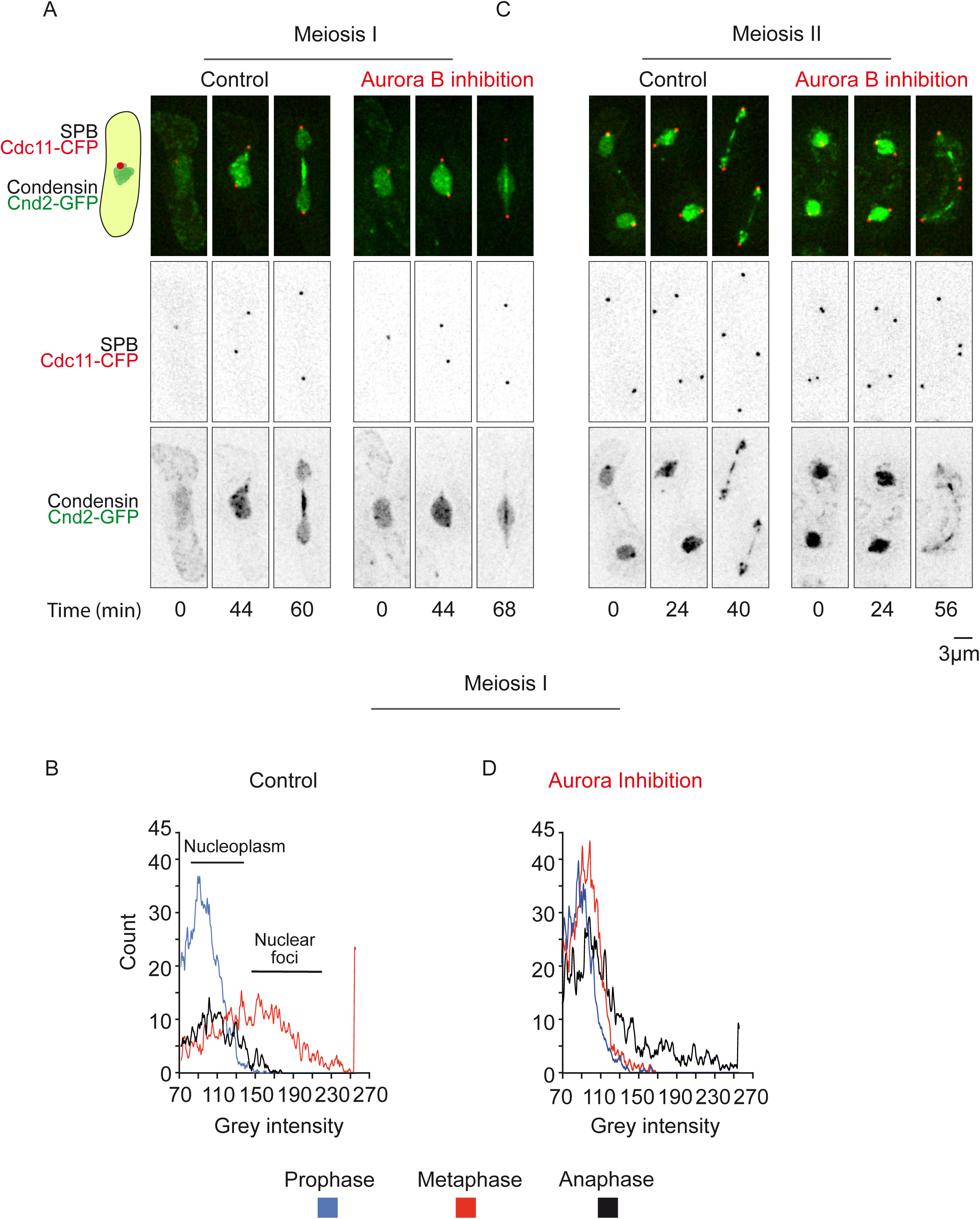
Aurora-dependent recruitment of condensin to chromosomes in both meiosis I and meiosis II. Condensin localization during meiotic progression is shown. SPBs are visualized via Cdc11-CFP (red) and condensin via Cnd2-GFP (green). **A**. Selected frames from a movie of *cnd2-GFP ark1-as3* cell undergoing meiosis I before (left panel) or after Aurora inhibition (right panel). **B**. Analysis of condensin recruitment using a histogram of pixel intensity. **C**. Selected frames from a movie of *cnd2-GFP ark1-as3* cell undergoing meiosis II without (left panel) or with (right panel) Aurora inhibition.

Similarly to what has been described in mitosis (Nakazawa et al., 2011; Tada et al., 2011), the presence of condensin foci in metaphase of meiosis I was clearly reduced after Aurora inhibition (Fig. 5A; right panel; movie 6). Consistent with this, the histogram analysis of foci pixel intensities following Aurora inhibition remained low from prophase to anaphase (Fig. 5D; right panel). A similar analysis of condensin localization in meiosis II again revealed the presence of several nuclear foci in metaphase. At anaphase II onset, condensin switches to chromosome arms and the spindle midzone (Fig. 5C; left panel; movie 7). Following Aurora inhibition, we observed the complete extinction of the chromosomal localization of condensin in anaphase II (Fig. 5C; right panel; movie 8) and only spindle midzone condensin signal remained present.

Thus, Aurora B controls formation of nuclear condensin foci in meiosis I and meiosis II as previously reported in mitosis. However, the function of Aurora B is only essential for chromosome arms and telomere separation in meiosis I but dispensable in meiosis II.

### Aurora B phosphorylation of condensin promotes chromosome arm/telomere separation in meiosis I

Previous studies in fission yeast have established that Cnd2 is a mitotic Aurora-B substrate (Nakazawa et al., 2011; Tada et al., 2011), and a Cnd2 phosphomimetic mutant (*cnd2-3E*) alleviates chromosome segregation defects in cells where Aurora B has been inactivated (Nakazawa et al., 2011; Reyes et al., 2015; Tada et al., 2011). We speculated that phosphorylation of Cnd2 by Aurora B may also be sufficient to promote telomere dissociation and chromosome arm separation in meiosis I. To address this point, we used *cnd2-3E ark1-as3* mutant cells that also expressed SPB (Cdc11-cfp, red) and telomere (Taz1-gfp, green) markers. We scored the number of Taz1 foci relative to spindle length in *cnd2-3E* cells after Aurora inhibition in meiosis I (Fig. 6A-C). As shown in Figure 6C, the *cnd2-3E* mutant completely bypassed telomere non-disjunction phenotype that arises from Aurora inhibition in meiosis I. Similarly and as expected, inhibiting Aurora in meiosis II in a *cnd2-3E* mutant had no impact on telomere dissociation (Fig. 6D-F). However, contrary to what has been observed in mitosis (Reyes et al., 2015), the *cnd2-3E* mutation did not alleviate the centromere attachment defects after Aurora B inactivation (Fig.S4). Indeed, *cnd2-3E* cells displayed numerous lagging and mis-segregated centromeres following Aurora inhibition (Fig.S4). Similarly, in meiosis II, while the effect of Aurora on telomere separation in this mutant was undistinguishable from control cells (Fig.6 D-F), Aurora’s role at centromeres, visualized through the presence of lagging and centromere mis-segregation, was independent of Cnd2 phosphorylation (Fig. S4).

**Figure 6:**
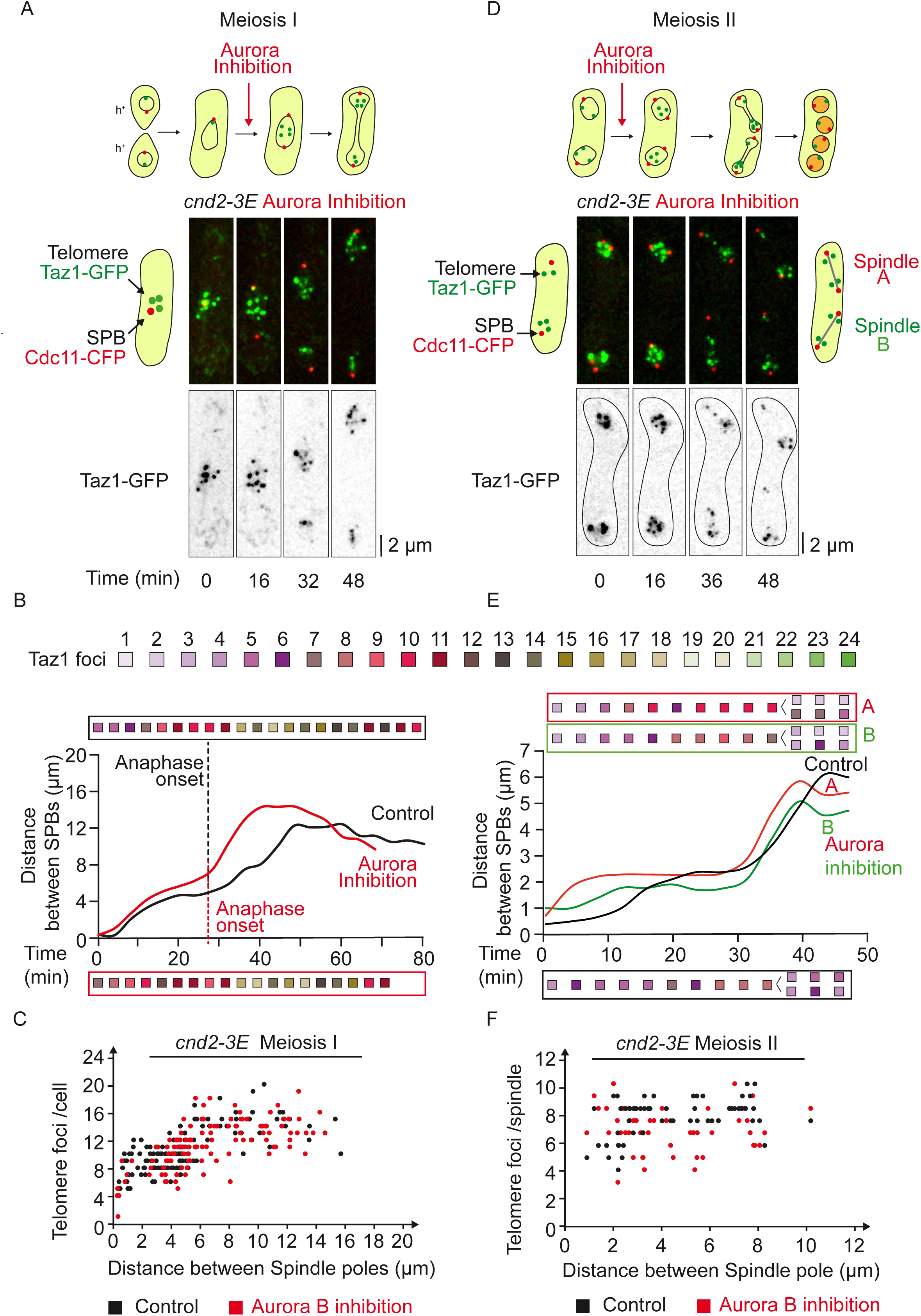
Condensin phosphomimetic mutant rescues telomere non-disjunction under Aurora inhibition in meiosis I. **A**. Selected frames from a movie of a live *cnd2-3E ark1-as3* cell undergoing meiosis I under Aurora inhibition. Telomeres are visualized via Taz1-GFP (green) and SPBs via Cdc11-CFP (red). **B**. Telomere foci dynamics during meiosis I in *cnd2-3E* cells. Analysis of the movie shown in A (Aurora Inhibition, red line and box), as well as the analysis of a control movie for *cnd2-3E* cell (Control, black line and box). The number of telomere foci is shown with a color code for each frame of the movie. **C**. Number of telomere foci according to the distance between SPB foci in *cnd2-3E ark1-as3* cells undergoing meiosis I. Data were measured from static images in control (n=137; black dots) and Aurora Inhibition (n=121; red dots). **D**. Selected frames from a movie of a live *cnd2-3E ark1-as3* cell under meiosis II-specific Aurora inhibition. Telomeres are visualized via Taz1-GFP and SPBs via Cdc11-CFP (red). **E**. Telomere foci dynamics in *cnd2-3E ark1-as3* cells under meiosis II-specific Aurora inhibition. Analysis of the movie shown in D; the upper and lower meiosis II spindles of the movie are defined as spindle A and spindle B respectively. The number of telomere foci is shown with a color code for each frame of the movie for both Spindle A (red box) and Spindle B (green box). Analysis of a control *cnd-3E ark1-as3* cell in meiosis II (Control, black line and box). After anaphase II onset, the number of telomere foci was counted separately in both sets of segregated chromosomes. **F**. Number of telomere foci according to the distance between SPB foci during meiosis II in *cnd2-3E ark1-as3* cells. Data was measured in meiosis II in control (n=6 movies; black dots) and after Aurora inhibition (n= 8 movies; red dots).

Together, these observations suggest that Aurora-B controls telomere and chromosome arm separation, through phosphorylation of the condensin Cnd2 subunit in meiosis I but not in meiosis II. As opposed to mitosis, Aurora’s role in correcting centromere attachment is largely but not totally independent of Cnd2 phosphorylation in meiosis I and II.

### Condensin is required for telomere and chromosome arm separation in meiosis I but not meiosis II

Our observations suggest that the activity of the condensin complex may be dispensable for telomere and chromosome arm separation in meiosis II. To verify this, we expressed SPB (Cdc11-CFP, red) and telomere (Taz1-GFP, green) markers cells bearing a conditional shut-off allele for the condensin subunit Cut14^Smc2^ (combining promoter repression with an auxin-inducible degron) (Kakui et al., 2017). To determine the effect of condensin degradation on chromosome arm and telomere separation, we analyzed the number of telomere foci with respect to spindle size in meiosis I and II (Fig.7 A-C). In meiosis I, even after full spindle elongation, the number of telomere foci was significantly reduced when condensin was degraded (Fig.7 A-B). This phenotype was similar to Aurora inhibition. However, in contrast to our observations following Aurora inhibition, condensin degradation led to few centromeric segregation defects (Fig.S4). Similarly, we addressed the role of Condensin in meiosis II by inducing condensin depletion (see material and methods; Fig.7 A-D). Interestingly, telomere and chromosome arm separation in anaphase II was not deeply impeded by condensin degradation (Fig.7A-C). To further analyze the role of Aurora and condensin in chromosome arm condensation, we measured the distance between SPBs of individual telomeric foci during meiosis II progression. As expected, condensin degradation and Aurora-inhibition caused chromosome arm condensation defects in anaphase of meiosis II as judged by the increase in SPB to telomere distances (Fig.7D; spindle size between 5 and 8 µm). Notably, the defect in chromosome arm condensation following Aurora inhibition was bypassed in the *cnd2-3E* phosphomimetic mutant (Fig.7D). However, at the end of meiosis II (post-elongation constant spindle size), chromosome arms were able to separate as judged by the decreasing distance between the SPBs and the telomeres (Fig.7D), although we could confirm that the localization of Cnd2-GFP was affected by Cut14 degradation both in meiosis I and II (Fig.8 A-B).

**Figure 7:**
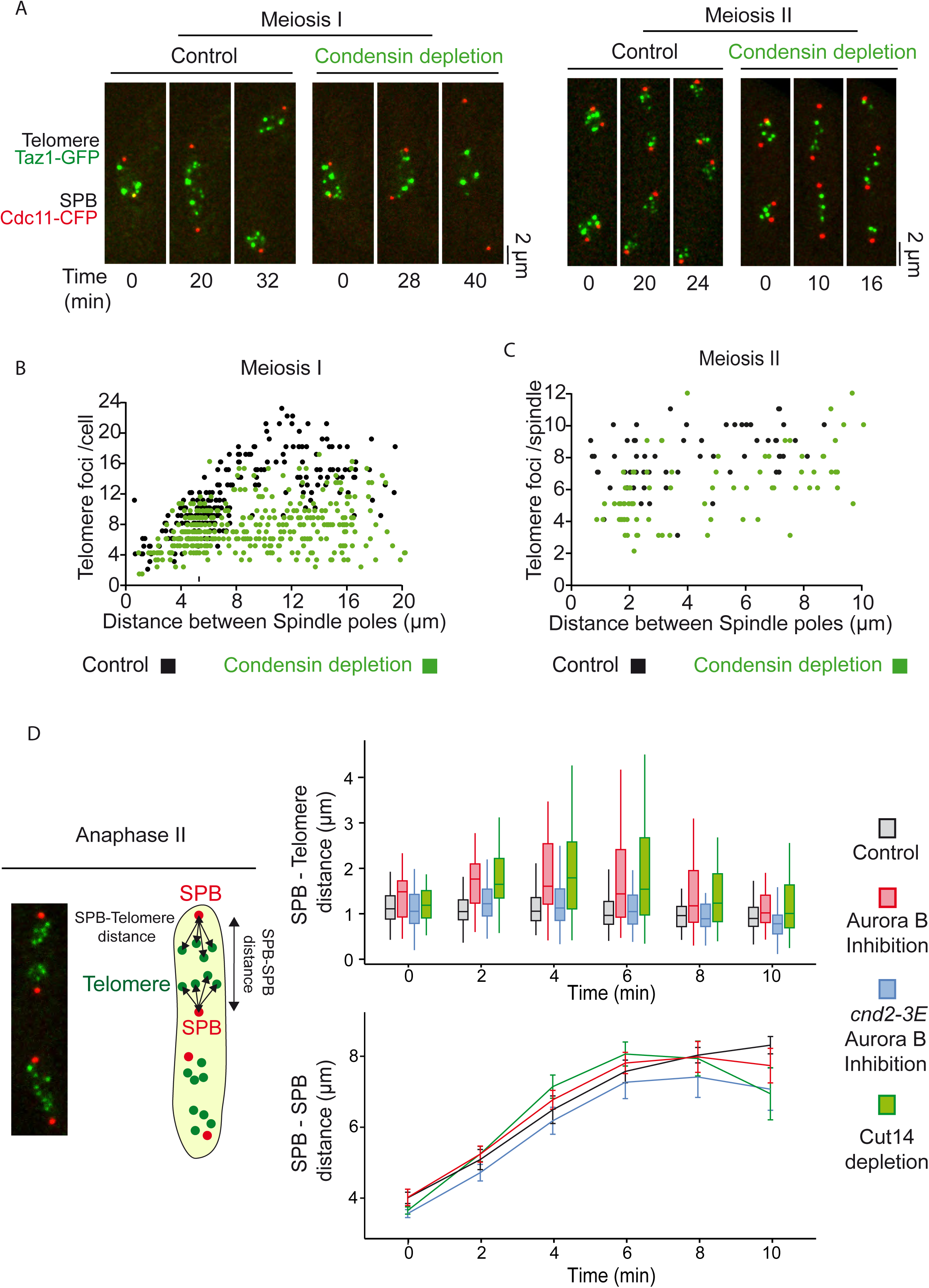
Telomere separation is achieved in meiosis II despite condensin depletion. **A**. Selected frames from movies of *cut14-shutoff* cells undergoing meiosis I (left panel) and meiosis II (right panel). Telomeres are visualized via Taz1-GFP (green) and SPBs are visualized via Cdc11-CFP (red). Condensin depletion is induced by addition of 0.5 mM of Auxin and 5μg/ml of thiamine. Image acquisition was started after 3h of induction for meiosis I and after 2h20 for meiosis II. **B**. Number of telomere foci according to the distance between SPB foci in meiosis I. Condensin depletion (green dots) is induced by addition of 0.5 mM of Auxin and 5μg/ml of thiamine. Control (black dots) corresponds to images of *cut14-so* cells grown in the absence of Auxin and thiamine. Data were measured from static images of control cells (n=226; black dots) and cells submitted to condensin depletion for 3h (n=337; green dots). **C**. Number of telomere foci according to the distance between SPB foci in meiosis II. Condensin depletion (green dots) is induced by addition of 0.5 mM of Auxin and 5 μg/ml of thiamine. Control (black dots) corresponds to *cut14-so* cells filmed in the absence of Auxin and thiamine. Data were measured on movies (n = 8 spindles, control cells, black dots; n = 8 spindles of cells depleted for condensin, green dots). **D**. Dynamics of telomere separation in meiosis II. Movies of cells in meiosis II were used to calculate the distances between each telomere foci and its nearest SPB focus (n = 8 spindles for each condition). Upper graph. The telomere to SPB distance is plotted according to the time from anaphase II onset which corresponds to t=0. The average corresponding SPB to SPB distance for the same movies are reported in the lower graph (mean +/− SD for each condition). Control (black boxes and line) are *ark1-as3* cells filmed in the absence of 1-NA-PP1; Aurora inhibition (red boxes and red line) is performed on *ark1-as3* cells, *cnd2-3E ark1-as3* cells (blue boxes and blue line) treated with 10µM 1-NA-PP1; Cut14 depletion (green boxes and green line) is performed on *cut14-so* cells by addition of 0.5 mM of Auxin and 5μg/ml of thiamine.

**Figure 8:**
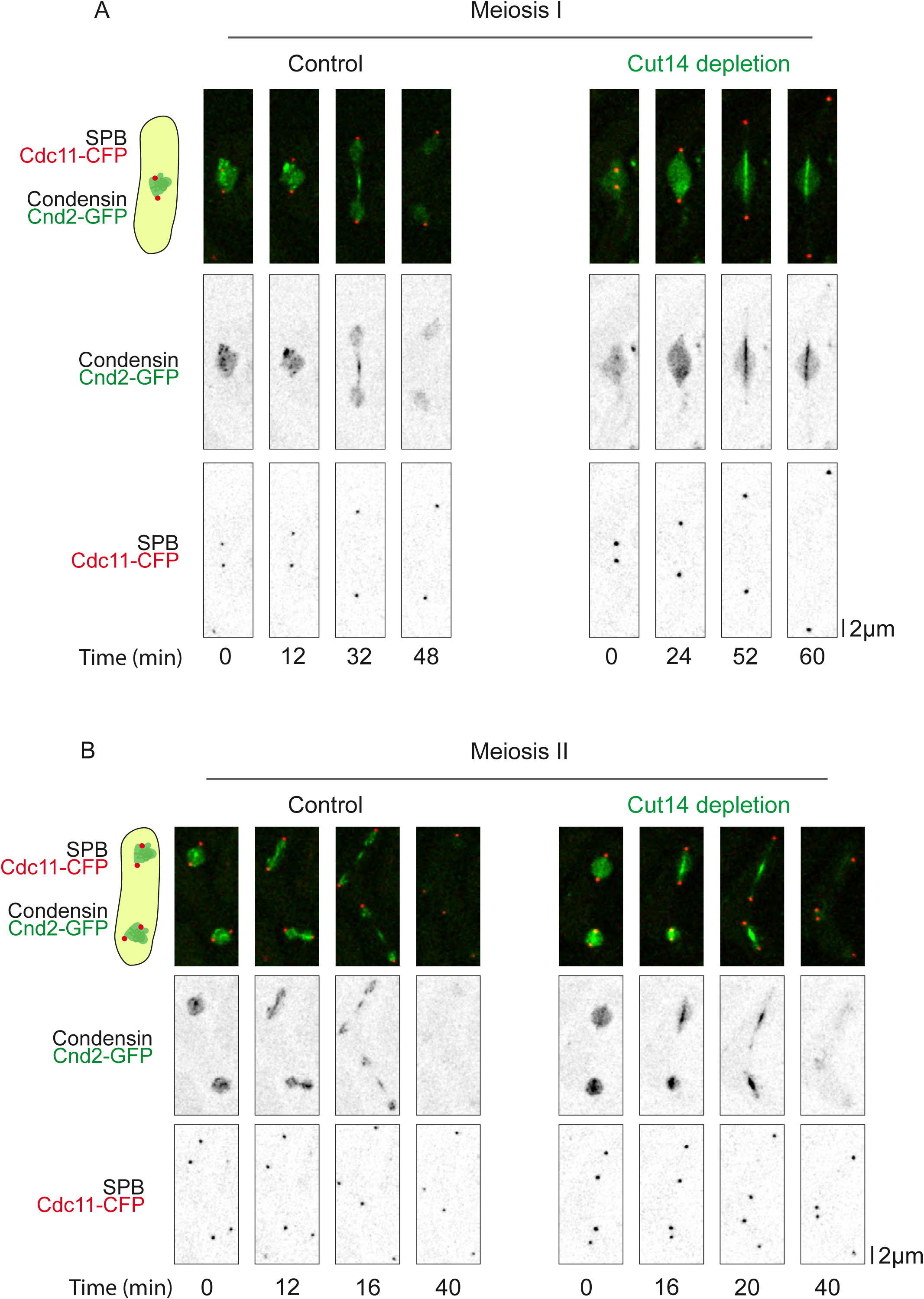
Cut14 depletion induces a loss of condensin localization to chromosomes. Selected frames from movies showing condensin localization upon Cut14 depletion in meiosis I (A) and meiosis II (B). SPBs are visualized via Cdc11-CFP (red) and condensin is visualized via Cnd2-GFP (green). WT (Control) and *cut14-so* (Cut14 depletion) cells were exposed to 0.5 mM of Auxin and 5μg/ml of thiamine for 3h in (A) and 2h20 in (B).

These observations confirm that telomere/chromosome arm separation requires condensin function in meiosis I but not in meiosis II. Furthermore, both in meiosis I and meiosis II, the importance of condensin in the control of KT-MT attachment remains limited.

Therefore, these results suggest the existence of an Aurora- and condensin-independent mechanism to segregate chromosome arms in meiosis II.

### Role of meiotic or mitotic cohesins in Aurora’s function at telomere/chromosome arms

It has been suggested that new cohesion on chromosome arms cannot be generated between meiosis I and II in the absence of de novo DNA replication (Toth et al., 1999; Uhlmann and Nasmyth, 1998; Watanabe and Nurse, 1999; Watanabe et al., 2001), we thought to test the role of cohesin in the chromosome arm/telomere phenotypes observed following Aurora inhibition. During meiosis I, cohesion is eliminated along chromosome arms to allow homologous segregation, but persists between sister centromeres moving towards the same spindle pole. The fission yeast cohesin protein Rec8 plays a key role in this process. Rec8 largely localizes all along chromosomes and persists at the centromeric regions throughout meiosis I until the anaphase of meiosis II (Watanabe and Nurse, 1999). To test whether Aurora B’s role in telomere and chromosome arm separation was dependent on the meiotic cohesin Rec8, we added the 1-NA-PP1 inhibitor to *rec8Δ ark1-as3* cells expressing SPBs (Cdc11-cfp, red) and telomere (Taz1-gfp, green) markers and followed cells progressing into meiosis I (Fig. 9A-B). Strikingly, even in the absence of Rec8, Aurora inhibition significantly reduced the number of Taz1 foci, as opposed to untreated *rec8Δ* cells (Fig. 9B). In agreement with this result, in *rec8^+^* cells, Rec8 remained solely associated in anaphase with the segregated centromeres (Fig. 9C) but no Rec8 foci co-localized with telomeres or chromosome arms following Aurora inhibition (Fig. 9C, Aurora inhibition) as previously reported (Hauf et al., 2007).

**Figure 9:**
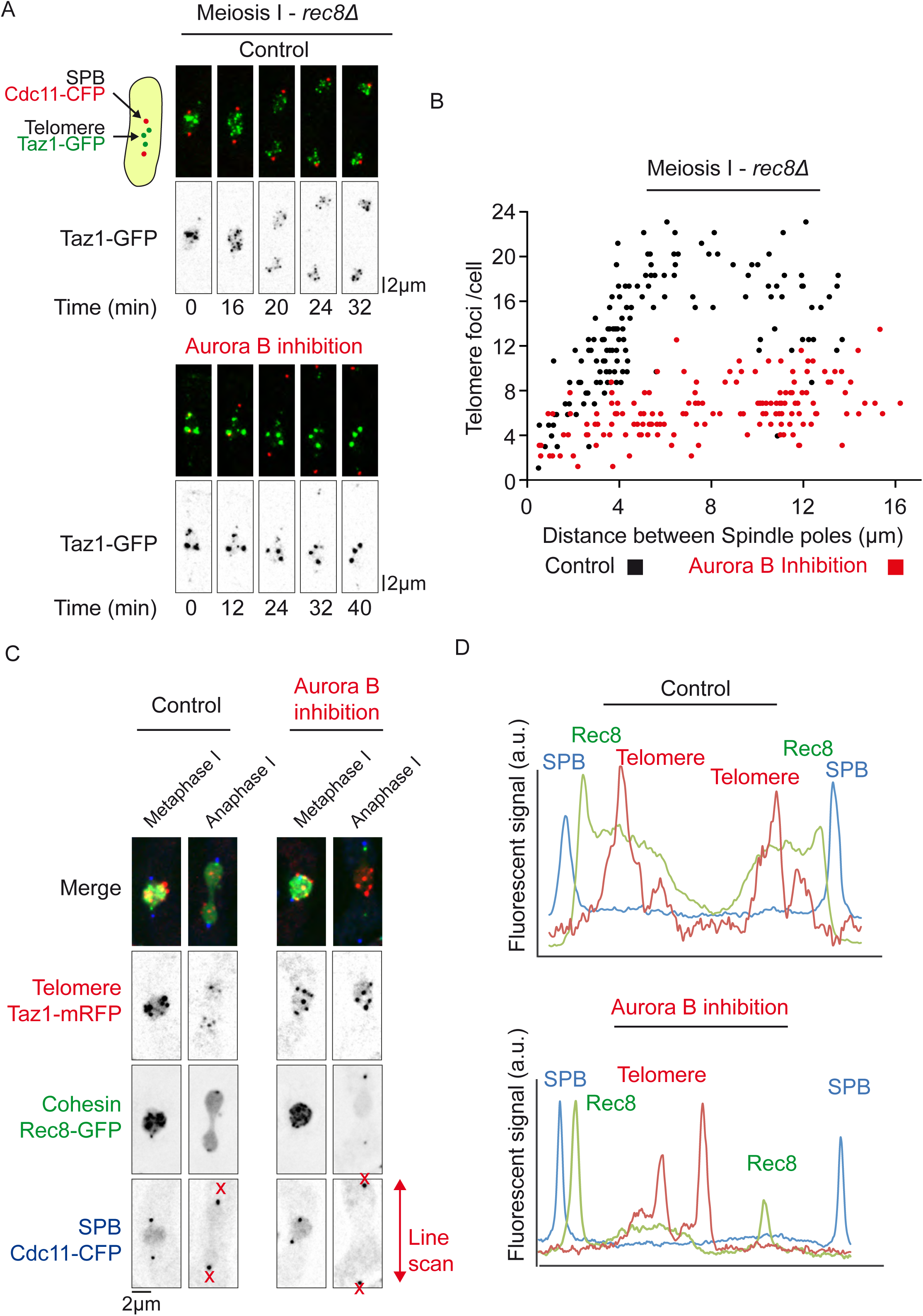
Aurora kinase is required for telomere separation in meiosis I independently of the meiotic cohesin Rec8. **A**. Selected frames from a movie of *rec8Δ ark1-as3* cell undergoing meiosis I. Telomeres are visualized via Taz1-GFP (green) and SPBs are visualized via Cdc11-CFP (red). **B**. Number of telomere foci according to the distance between SPB foci in *rec8Δ ark1-as3* cells undergoing meiosis I. Data was measured from static images of *rec8Δ ark1-as3* cells (n=142, black dots) and *rec8Δ ark1-as3* cells under Aurora Inhibition (n=148, red dots). **C**. Meiotic cohesin Rec8 localization during meiosis I. Static images of live cells representative of metaphase I and anaphase I. Telomeres are visualized via Taz1-mRFP (red), meiotic cohesin via Rec8-GFP (green) and SPBs via Cdc11-CFP (blue). D. Line scans analysis of the cells shown in C, for control (top panel) and Aurora inhibition (lower panel).

In contrast to the behavior of Rec8 in meiotic cells, the mitotic cohesin Rad21 is found predominantly at the leading edge of the nuclei undergoing prophase oscillations during meiosis I, just where the telomeres are clustered (Watanabe and Nurse, 1999; Yokobayashi et al., 2003). In addition, it has been shown that the equational segregation of chromosomes in meiosis I in the absence of Rec8 depends on Rad21, which is then able to locate to the centromeres (Yokobayashi et al., 2003). In light of these findings, we analysed the localization of Rad21 in wild type or Rec8 deleted cells (Fig.S5). In addition to the fraction co-localizing with the nucleolus/telomere region of prophase nuclei (Fig.S5A, control), we detected Rad21 foci at the telomere/nucleolar region of metaphase cells (Fig.S5A, control). These foci rapidly disappeared at the metaphase to anaphase transition as previously seen during mitosis (Reyes et al., 2015). This pattern of Rad21 localization was also observed at the metaphase to anaphase transition when Rec8 was deleted (Fig.S5C, control) but as previously described, Rad21 also co-localized strongly to the centromere cluster in prophase of meiosis I (Fig.5C, control). Interestingly, a small fraction of Rad21 remained at nucleoli and telomeres at anaphase when Aurora kinase was inactivated, in both control and *rec8Δ* cells (Fig.S5A-D, aurora inhibition).

Together, these results demonstrate a role for Aurora at telomeres/chromosome arms in meiosis I, independently of the meiotic cohesin Rec8 and also suggest the existence of an Aurora- and condensin-independent mechanism of chromosome segregation in meiosis II.

## Discussion

Our study reveals that the physical association of telomeres is tightly regulated during meiotic division. Interestingly, we observe both similarities and differences with mitosis in the dynamics of telomere separation. During mitosis, telomere foci first dissociate from the nuclear envelope (Fujita et al., 2012) and then undergo separation in two discrete steps, both dependent on Aurora activity. Telomere dispersion occurs during metaphase, prior to chromosome segregation, whilst sister chromatid telomere disjunction is achieved during mid-anaphase (Reyes et al., 2015). We previously proposed that the dissociation of the Shelterin complex in early mitosis initiates telomere dispersion, promotes condensin loading and participates in chromosome arm separation. Similarly, in meiosis I, our study demonstrates that telomere separation occurs in two steps, dispersion in metaphase and disjunction during anaphase I. We further show that both steps are dependent on Aurora B kinase activity. Thus the mechanisms underlying telomere separation might be similar during mitosis and meiosis I (Fig.10).

**Figure 10:**
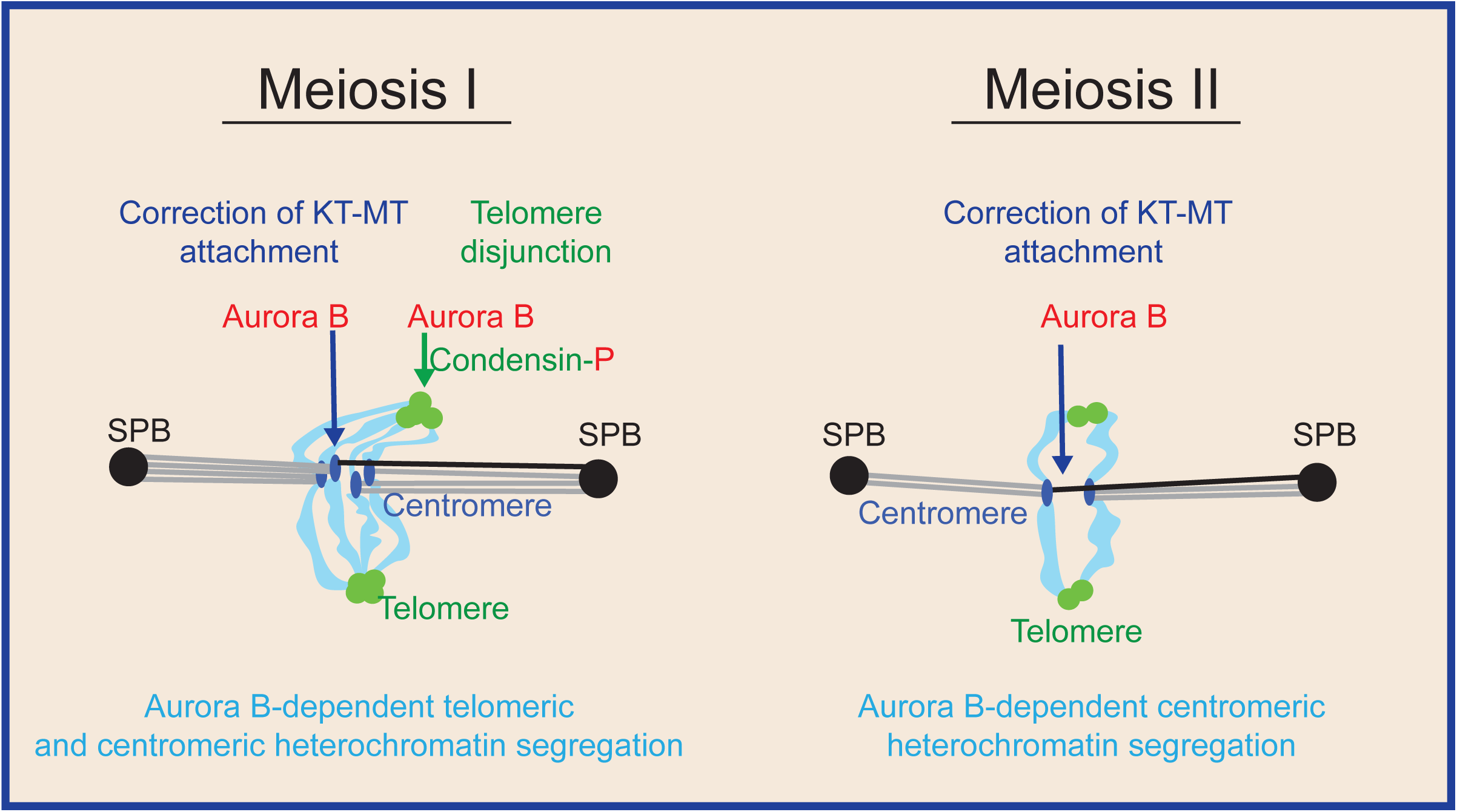
Model summarizing the respective roles of Aurora and condensin in centromeric, telomeric or chromosome arms segregation in meiosis I and II. Schematic representation illustrating the regulation of telomere, chromosome arms, centromere dissociation throughout meiosis.

In meiosis I, we show that Aurora B spatially targets distinct chromosomal domains (i.e. telomeres and centromeres) to control KT-MT attachment and telomere/chromosome arm separation. Previous studies have reported that the Shugoshin protein Sgo2 (Hauf et al., 2007; Vanoosthuyse et al., 2007) or the cohesin Rad21 (Morishita et al., 2001) were required for the proper recruitment of Aurora at centromeres. However, what controls the loading of Aurora at fission yeast telomeres remains to be clarified since many factors seem to contribute to its localization. In mitosis, Aurora B promotes telomere and chromosome arm separation by phosphorylating and recruiting the condensin subunit Cnd2 to chromosomes (Nakazawa et al., 2011; Reyes et al., 2015; Tada et al., 2011). Yet, the role of Aurora in correction of merotelic attachment is only partially dependent on the phosphorylation of Cnd2 condensin subunit since a phosphomimetic mutant of Cnd2 (*cnd2-3E*) only moderately bypasses chromosome arm separation defects following Aurora inhibition (Nakazawa et al., 2011; Reyes et al., 2015; Tada et al., 2011).

In the present study, we went further in characterizing the role of condensin in KT-MT attachment and chromosome arm separation during meiosis. Through specific depletion of condensin in meiosis I, we demonstrate that faithful KT-MT attachment is largely independent of condensin. Instead, condensin plays an essential role in telomere/chromosome arm separation. A phosphomimetic mutant of *cnd2* (*cnd2-3E*) is able to bypass telomere/chromosome arm separation defects following Aurora inhibition. But again, in meiosis I, the function of Aurora in KT-MT attachment is completely independent of Cnd2 phosphorylation (Fig.10). How does condensin trigger chromosome arm-telomere separation? Similarly to the role of spindle elongation forces in anaphase error correction (Choi and McCollum, 2012; Cimini et al., 2004; Courtheoux et al., 2009; Gay et al., 2012), we speculate that condensin may control telomere separation by mechanically pulling on chromosome arms.

During telophase of meiosis I, the spindle disassembles, telomeres reform clusters at the nuclear periphery prior to the onset of meiosis II. Thus, the reclustering of telomeres at the nuclear periphery is independent of *de novo* DNA replication. Finally, in meiosis II, telomeres separate in a single step during metaphase, prior to chromosome segregation and independently of Aurora activity. In this respect, the dynamics of telomere separation in meiosis II differs from meiosis I and mitosis. In meiosis II, Aurora B only localizes to centromeres, which allowed us to separate the functions of the CPC at those chromosomal loci. Our observation that condensin is largely dispensable in meiosis II for telomere/chromosome arm separation, might reflect lower opposing forces to spindle elongation in meiosis II as opposed to mitosis or meiosis I. However, we found no clear differences in the rate of spindle elongation in mitosis/meiosis I or meiosis II cells arguing against this hypothesis. Alternatively, a chromatin compaction mechanism associated with spore formation might compensate for reduced condensin activity during meiosis II.

A major difference between meiosis I and II is the specific presence in meiosis II of a phenomenon called ‘‘virtual’’ NEBD (vNEBD) (Arai et al., 2010; Asakawa et al., 2010), a key event for *S. pombe* meiosis (Asakawa et al., 2016; Flor-Parra et al., 2018). Although the nuclear envelope and the Nuclear Pore Complexes remain assembled throughout the entire meiotic division, nuclear proteins are able to diffuse transiently into the cytoplasm at the onset of anaphase B of meiosis II. Consequently, nuclear and cytoplasmic molecules appear to be mixed in a process resembling an open mitosis. This phenomenon certainly accounts for some of our observations such as the disappearance of condensin signal from the nucleus following Aurora inhibition in meiosis II. Hence, it remains possible that the diffusion of several nuclear proteins in meiosis II participates to this non-canonical chromosome segregation event. A second difference between meiosis I or II is the presence in meiosis II of the forespore membrane, which encapsulates the nucleus and produces forces for cell division (Akera et al., 2012). Markedly, it has been shown that interpolar MTs were dispensable in meiosis II for spindle pole separation (Akera et al., 2012). Finally, it is formally possible that the lack of DNA replication prior meiosis II might change the requirement for condensin activity during the subsequent division. New cohesion on chromosome arms cannot be generated between meiosis I and II in the absence of de novo DNA replication (Toth et al., 1999; Uhlmann and Nasmyth, 1998; Watanabe et al., 2001). Meiosis II chromosomes may thus lack some specific features dependent on cohesins. In mitosis, the dispersal of Rad21 from telomeres relies upon Aurora B (Reyes et al., 2015). It has also been reported that cohesin participates in subtelomeric heterochromatin maintenance in fission yeast probably by acting locally on Swi6/HP1 binding in the subtelomeric region (Dheur et al., 2011). However, we found that the role of Aurora-B on telomere/chromosome arm separation in meiosis I was independent of the key meiotic cohesin, Rec8. In addition, Rec8 was not enriched at telomeres before or after Aurora inhibition. Rec8 was first identified in a genetic screen for mutants showing a decrease in recombination at the ade6 locus (Ponticelli and Smith, 1989) and several features of meiotic chromosomes were shown to be disrupted in *rec8Δ* cells, including the formation of chiasmata (Ellermeier and Smith, 2005; Molnar et al., 1995; Watanabe and Nurse, 1999). Our results therefore suggest that the telomere dissociation pathway in meiosis I operates independently of chiasma resolution.

Conversely and similarly to our observations in mitosis, we were able to detect a subfraction of Rad21 cohesin that remained associated with telomeres/nucleolus following Aurora inhibition. Thus, the presence of this subfraction of cohesin during anaphase may be sufficient to maintain telomere entanglements in meiosis I. If this is the case, the role of Aurora kinase at fission yeast telomeres resembles the function of tankyrase 1, since the resolution of sister chromatid cohesion between telomeres in human cells also occurs in early mitosis but has a specific requirement for the poly(ADP-ribose) polymerase (PARP) tankyrase 1 (Bisht et al., 2013; Dynek and Smith, 2004).

In conclusion, it is often mentioned in the scientific literature that chromosome segregation in mitosis or meiosis II requires similar mechanisms since both processes involve the separation of sister-chromatids to the two daughter cells. Our study challenges this view by revealing fundamental differences existing between mitosis and meiosis II in the control of chromosome segregation. Unlike mitosis or even meiosis I, chromosome segregation during meiosis II is largely independent of Aurora kinase and condensin, arguing that it most likely involves hitherto ignored non-canonical mechanisms.

## METHODS

### S. pombe strains and culture

Media, growth, maintenance of strains and genetic methods were as described (Moreno et al., 1991). All strains used in this study are listed in supplementary table 1. All featured alleles are created from original strains, referenced and described in supplementary table 1. Crosses for strain generation were performed by tetrad dissection and selected either through drug resistance, prototrophy for specific amino acids or visual selection for fluorescent markers. The *cnd2-3E* allele was selected in *ark1-as3* strains by testing the resistance to Aurora inhibition with 1µM 1-NA-PP1. Strains were grown on YES solid media at 25°C. Strains carrying Cut14 under the expression of the nmt81 promoter were grown on EMM+N solid media at 25°C. To induce meiosis, h+ and h-strains were mixed (or h90 alone) and spread on EMM-N solid media at 25°C overnight.

### Live-cell microscopy

To increase the density of meiotic cells prior to filming, crosses were suspended in liquid EMM media and aggregates of meiotic cells were left to sediment at the bottom of the tube. Supernatant was discarded and cells were resuspended three times. Meiotic cells were spread on a plug of EMM filled with 2% agarose in an imaging chamber (CoverWell imaging chamber 22mm x 1.7mm Grace Bio-Labs). The imaging chamber was sealed with a 22×22 glass coverslip.

Images were acquired with a CCD CoolSNAP HQ2 camera (Roper Scientific) fitted to a DM6000 upright microscope (Leica) with a x100 1.40NA objective or a CCD Retiga R6 camera (QImaging) fitted to a DM6B upright microscope with a x100 1.44NA objective, using Metamorph as a software. Z-stacks of 9 planes spaced by 0.7µm were typically acquired at 4min intervals (2min for anaphase II analysis in Fig. 7D) for the entire duration of meiosis I and II at 25°C.

For CENII segregation analysis in Fig.S2 and Fig.S4 as well as for spindle elongation measurements in Fig. S1C and Fig. S1D, images were acquired with a Neo sCMOS camera (Andor) fitted to an Eclipse Ti inverted microscope (Nikon) with a x60 1.40NA objective using MicroManager as a software at 2min intervals for 1h at 25°C.

### Aurora kinase inhibition and condensin depletion

The *ark1-as3* analogue sensitive allele (Koch et al., 2011) was used to perform Ark1 inhibition. To inhibit Aurora in meiosis I or II, the imaging media was supplemented with 10µM 1-NA-PP1 (Euromedex). However, to specifically inhibit Aurora kinase in meiosis II, image acquisition started exactly 10min after the spreading of cells on the imaging media containing 1-NA-PP1 and only cells starting meiosis II were filmed. This protocol allows imaging of cells that underwent normal chromosome segregation in meiosis I.

condensin depletion was performed using the *cut14-so* strain previously described (Kakui et al., 2017) that combines transcriptional repression of *cut14* under the nmt81 thiamine-repressible promoter and Cut14 depletion using an Auxin-Inducible Degron (AID) system. To induce condensin depletion, the imaging media was supplemented with 0.5mM of 1-*Naphthaleneacetic acid* (Sigma) and 5μg/ml of thiamine. For condensin depletion in meiosis I, image acquisition was started 3 hours after spreading the cells on the imaging media containing Naphthaleneacetic acid. For condensin depletion in meiosis II, image acquisition was started between 2 to 2h30 after spreading cells on the imaging media containing Naphthaleneacetic acid since longer induction caused meiosis I spindle collapse during the incubation period.

### Image processing and analysis

All image processing and analysis was performed using ImageJ. Deconvolution of Z-stacks and PSF generation were performed using the DeconvolutionLab and PSF Generator ImageJ plugins. Deconvolved stacks were combined using maximum intensity Z-projections. If needed, sample drifting over time was corrected with the MultiStackReg ImageJ plugin. Figures were assembled with the R ggplot2 package, ImageJ, Microsoft Excel and Adobe Illustrator. The number of telomere and Ark1 foci were assessed manually. Distances shown in Fig.S1B were measured on non-projected Z-planes. We measured the distances between the center of the telomere focus to the center of the nuclear envelope fluorescent signal. Distances in other figures were measured on Z-projections between the centers of telomere to SPB foci. In Fig.S2 and Fig.S4, lagging centromeres were defined as CENII foci that don’t co-localize with an SPB during anaphase I or II. Unequal centromere segregation was defined as a single CENII signal observed on a single SPB during anaphase I or II. In meiosis II, the two spindles were considered independently of one another. Spindle elongation profiles shown in Fig.S1C and Fig.S1D were measured on the movies of CENII segregation analysis. Anaphase onset was manually annotated and defined as the sharp increase in spindle elongation speed and spindle elongation curves were all aligned at anaphase onset.

For the analysis of condensin particle size, we performed an histogram of pixel intensities using imageJ.

## Online supplemental Material

Fig. S1 illustrates the clustering and the anchoring of telomeres throughout meiotic divisions.

Fig. S2 shows that Aurora exclusively co-localizes with telomeres during metaphase I.

Fig. S3 describes that Aurora kinase is required for proper attachment of KTs to the spindle during meiosis I and II.

Fig. S4 reveals that condensin is not required for KT-MT attachment in meiosis I or meiosis II.

Fig. S5 shows that a subfraction of Rad21 remains associated to telomere/nucleolus loci

Table S1 shows the strains used in this study.

## Acknowledgment

We would like to thank J. Cooper, J.P. Javerzat, M. Yanagida, for supplying strains; S. Hauf for supplying the *ark1-as3* mutant and Y. Watanabe for the *cnd2-3E* mutant; J.P. Javerzat, Dean Dawson for helpful discussions. J.B. was supported by a University MRT and Fondation pour la Recherche Medicale (FRM) fellowships. This work was funded by the ANR-16-CE12-0015 -TeloMito-2016 and l’Association de la Recherche sur le Cancer (ARC), fonctionnement, 2016. The microscopy equipment was funded by the CNRS and l’Association de la Recherche sur le Cancer (ARC). The authors declare no competing financial interests.

## ONLINE SUPPLEMENTAL MATERIAL

Supplemental data includes the strain list, five figures and 8 movies.

**Supplemental Figure S1:**
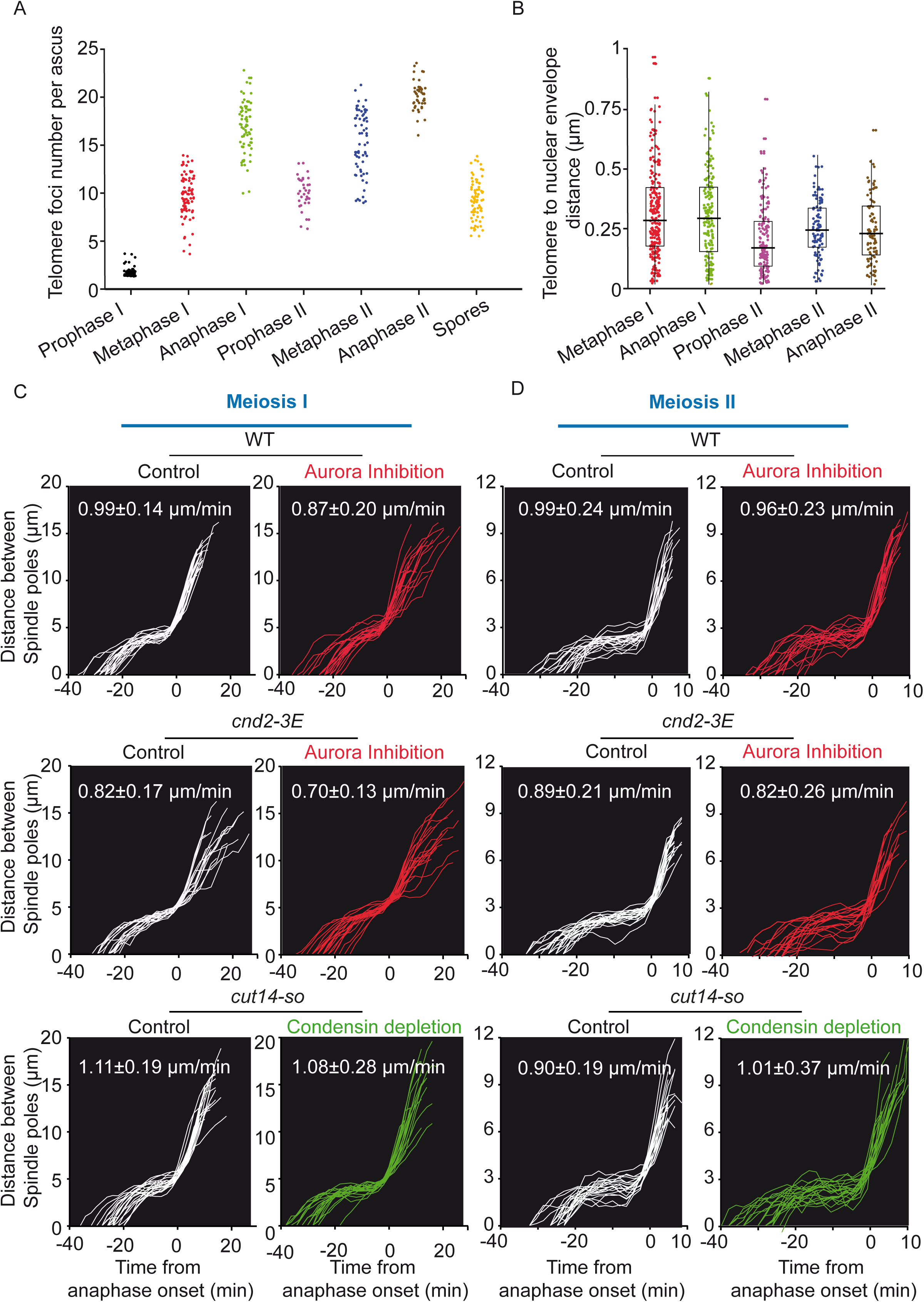
Telomere clustering, envelope anchoring and spindle elongation dynamics throughout meiotic divisions. **A**. State of telomere clustering throughout meiosis. Telomere foci number (Taz1-GFP) was assessed on static images of cells for each meiotic phase. Telomere foci number is reported per ascus. Thus, telomere foci number for both meiosis II and the 4 spores are therefore counted and reported together (n = 64 for Prophase I; n = 82 for Metaphase I; n = 70 for Anaphase I; n = 35 for Prophase II; n = 66 for Metaphase II; n = 45 for Anaphase II; n = 76 for Spores). **B**. Telomere positioning at the nuclear periphery during meiosis. Distances from telomere foci (Taz1-mRFP) to the nuclear envelope (Amo1-GFP) were measured on static images of live cells and reported for individual telomeres from Metaphase I to Anaphase II (n= 237 foci from 49 cells for Metaphase I; n = 179 foci from 31 cells for Anaphase I; n=155 foci from 34 cells for Prophase II; n = 104 foci from 17 cells for Metaphase II and n = 95 foci from 13 cells for Anaphase II). **C-D**. Spindle elongation dynamics in *ark1-as3*, *cnd2-3E ark1-as3* and *cut14-so* cells in meiosis I (C) and meiosis II (D). Distance between SPBs was measured on movies of live cells and is reported for individual cells according to time (t=0 corresponds to Anaphase I or II onset). For each condition more than 19 movies are plotted apart for *cnd2-3E* control in meiosis I where 16 movies are analyzed. Maximum spindle elongation speed during anaphase B of meiosis I or II is reported in the graph (mean +/− SD in µm/min). Aurora inhibition (red curves) was performed by addition of 10µM 1-NA-PP1 10min before the start of image acquisition. Condensin depletion (green curves) was performed by addition of 0.5mM of 1-*Naphthaleneacetic acid* (Auxin) and 5μg/ml of thiamine for 3h before the start of image acquisition. For condensin depletion in meiosis II, only cells that did not suffer obvious meiosis I defects were considered.

**Supplemental Figure S2:**
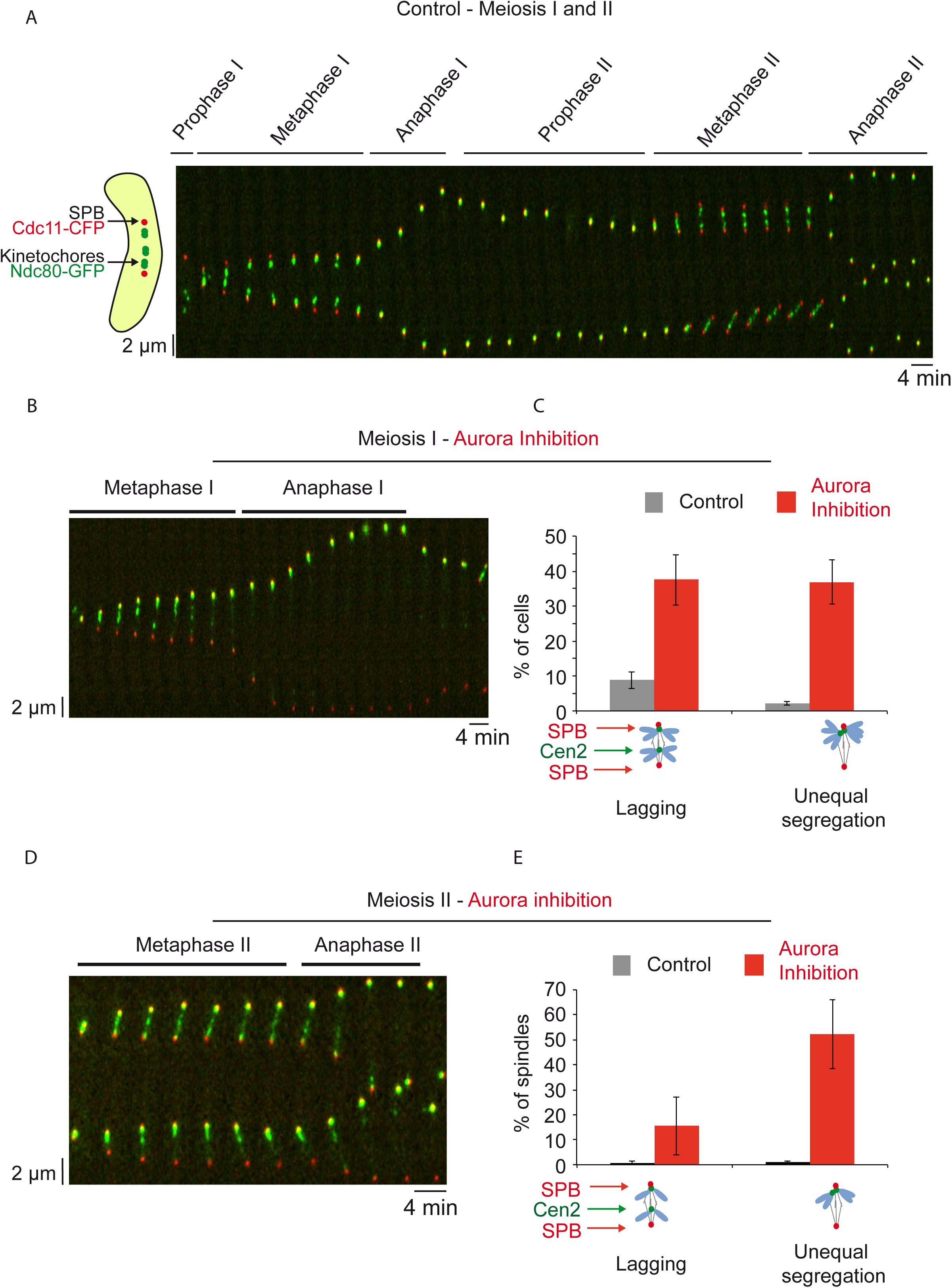
Aurora kinase is required for proper attachment of KTs to the spindle during both reductional and equational meiotic divisions. **A-D**. Timelapse imaging of KTs throughout meiotic divisions. Kymographs from movies of live *ark1-as3* cells. KTs are visualized via Ndc80-GFP (green) and SPBs are visualized via Cdc11-CFP (red). **A**. Control cell undergoing meiosis I and meiosis II successively. **B**. Cell undergoing meiosis I under Aurora inhibition. Aurora inhibition was performed by addition of 10µM 1-NA-PP1. **C**. Percentage of KT-MT attachment defects in the presence or absence of Aurora inhibition in meiosis I. Segregation pattern of centromere 2 was assessed in movies of live *ark1-as3* cells with homozygous tagging of a centromere 2 proximal loci (*lacO* insertions and expression of LacI-GFP). Lagging centromeres were defined as cells in which at least one centromere 2 focus does not colocalize with SPBs during anaphase I. Unequal segregation of centromere 2 was defined as cells in which only one focus of centromere 2 is observed and co-localizes with a SPB during anaphase I (Control, n = 411 cells from 3 independent experiments; Aurora inhibition, n = 363 cells from 4 independent experiments. **D**. Cell undergoing meiosis II under Aurora inhibition. Aurora inhibition was performed by addition of 10µM 1-NA-PP1, 10min before the start of image acquisition. **E**. Percentage of KT-MT attachment defects in the presence or absence of Aurora inhibition in meiosis II. The segregation pattern of centromere 2 in meiosis II was assessed in movies of live *ark1-as3* cells as explained above (Control, n=382 spindles from 3 independent experiments; Aurora inhibition, n = 158 spindles from 4 independent experiments).

**Supplemental Figure S3:**
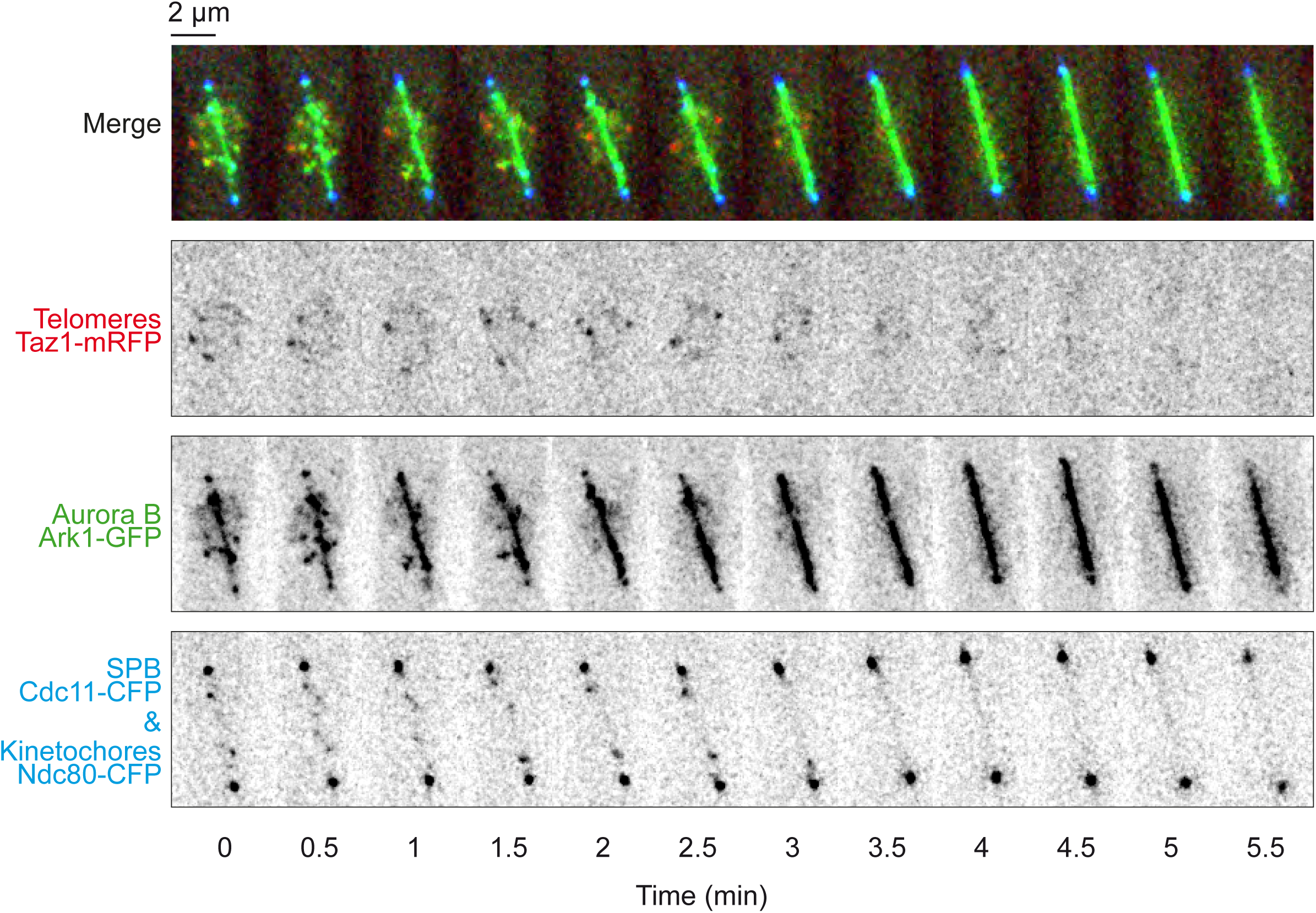
Aurora exclusively co-localizes with telomeres during metaphase I. Time lapse imaging of Metaphase I to Anaphase I transition in a live cell. Telomeres are visualized via Taz1-mRFP (red), Aurora is visualized via Ark1-GFP (green), SPBs are visualized via Cdc11-CFP (blue) and KTs are visualized via Ndc80-CFP (blue). Frames were acquired at an interval of 30s.

**Supplemental Figure S4:**
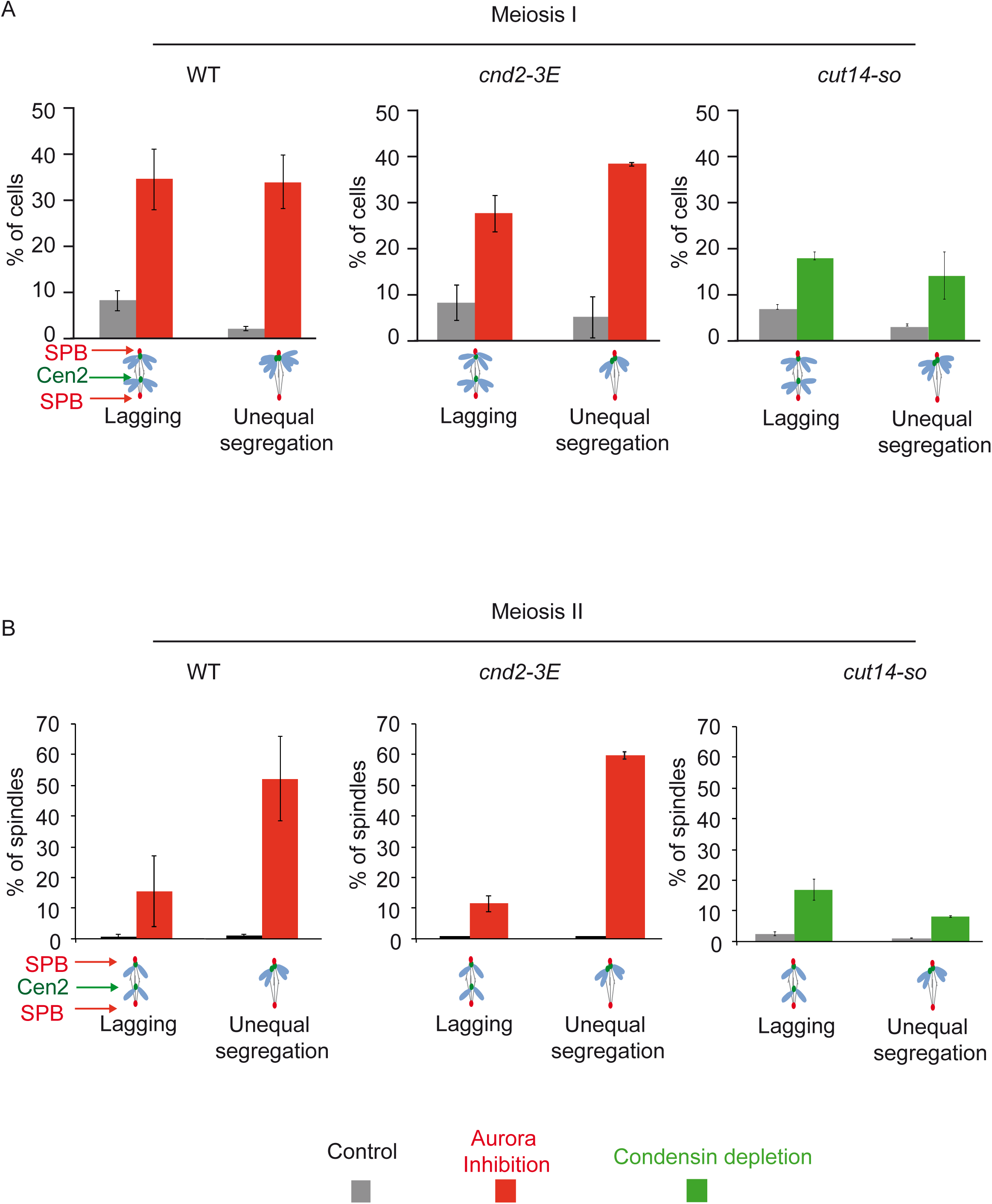
Condensin does not play a critical role in KT-MT attachment in meiosis I or meiosis II. The segregation pattern of centromere 2 was assessed in meiosis I (A) or meiosis 2 (B) in movies of live *cnd2-3E ark1-as3* and *cut14-so* cells with homozygous tagging of a centromere 2 proximal loci (*lacO* insertions and expression of LacI-GFP). Lagging centromeres were defined as cells in which at least one centromere 2 focus does not co-localize with SPBs in anaphase I or II. Unequal segregation of centromere 2 was defined as cells in which only one focus of centromere 2 is observed and co-localizes with a SPB in anaphase I or II. Aurora inhibition (red) was performed by addition of 10µM 1-NA-PP1, 10min before the start of image acquisition. Condensin depletion (green) was performed by addition of 0.5mM of 1-*Naphthaleneacetic acid* (Auxin) and 5μg/ml of thiamine for 3h before the beginning of image acquisition. For condensin depletion in meiosis II, only cells that did not suffer obvious meiosis I defects were considered. For comparison purpose, data from figure S2C and figure S2E were reported in WT panels. **A**. n = 118 cells from 2 independent experiments for *cnd2-3E* control; n = 226 cells from 2 independent experiments for *cnd2-3E* with Aurora inhibition; n = 117 cells from 2 independent experiments for *cut14-so* control and n = 111 cells from 2 independent experiments for condensin depletion with *cut14-so*. **B**. n = 221 spindles from 2 independent experiments for *cnd2-3E* control; n = 258 spindles from 2 independent experiments for *cnd2-3E* with Aurora inhibition; n = 222 spindles from 2 independent experiments for *cut14-so* control and n = 194 from 2 independent experiments for condensin depletion in *cut14-so*.

**Supplemental Figure S5:**
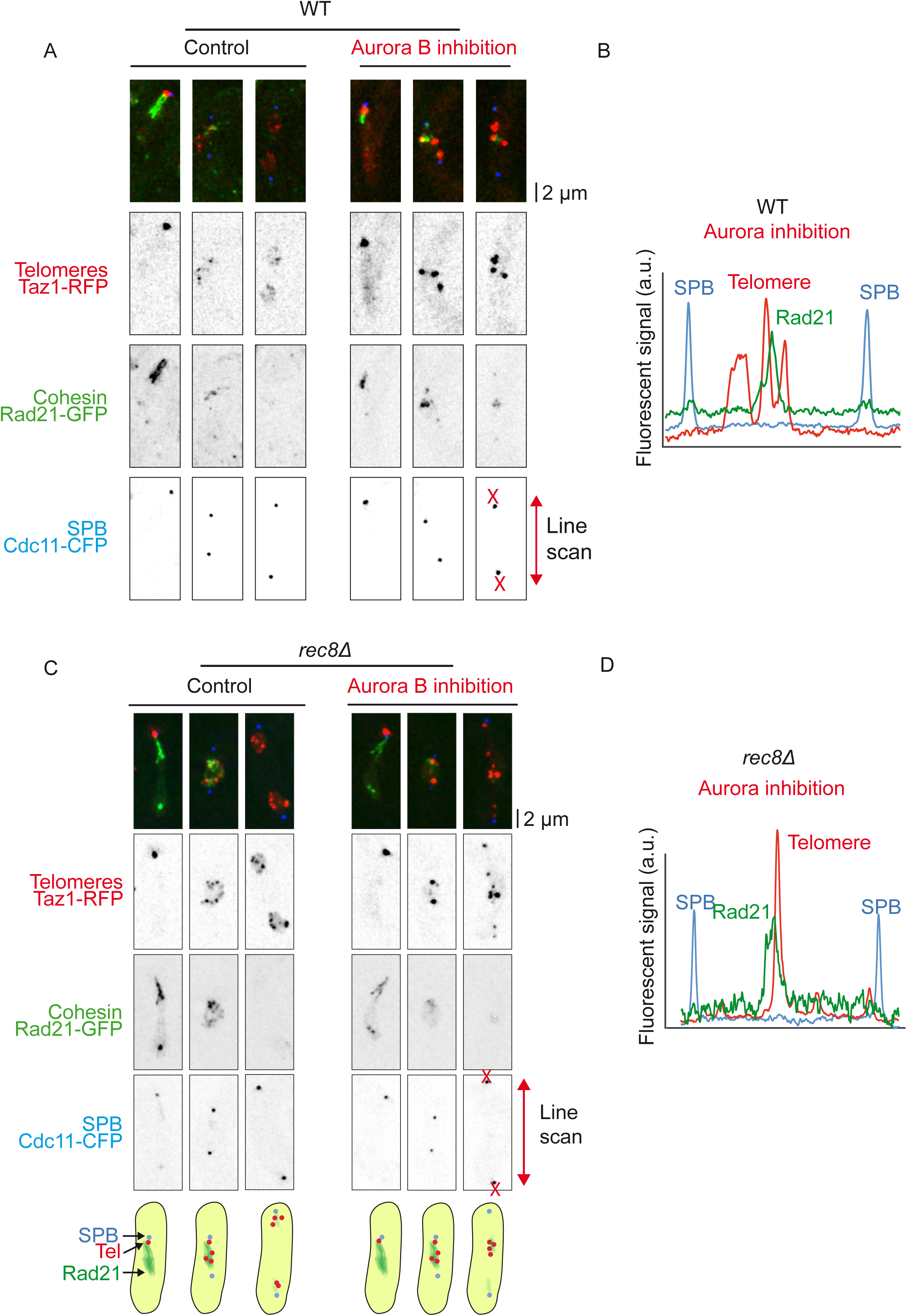
Aurora B controls the dissociation of a subfraction of cohesin Rad21 from telomeres/nucleolus at anaphase of meiosis I. **A**. Static images of live *ark1-as3* cells undergoing meiosis I. Telomeres are visualized via Taz1-mRFP (red), Cohesin is visualized via Rad21-GFP (green) and SPBs are visualized via Cdc11-CFP (blue). Aurora inhibition was performed by addition of 10µM 1-NA-PP1. **B**. Line scan analysis of the cell shown in A. Note the presence of a weak subfraction of Rad21 co-localizing with nucleolus/telomeres. **C**. Static images of live *rec8Δ ark1-as3* cells undergoing meiosis I. Telomeres are visualized via Taz1-mRFP (red), Cohesin is visualized via Rad21-GFP (green) and SPBs are visualized via Cdc11-CFP (blue). Aurora inhibition was performed by addition of 10µM 1-NA-PP1. **D**. Line scan analysis of the cell shown in C. Note the presence of a subfraction of Rad21 co-localizing with nucleolus/telomeres.

**Video 1: Fluorescent time-lapse imaging of telomere dynamics (Taz1-gfp) during meiosis I in fission yeast.**

Movie of a wild type cell expressing Taz1-gfp (telomeres, green) and Cdc11-cfp (SPBs, red).

**Video 2: Fluorescent time-lapse imaging of telomere dynamics (Taz1-gfp) during meiosis I following Aurora B inhibition.**

Movie of a wild type cell expressing Taz1-gfp (telomeres, green) and Cdc11-cfp (SPBs, red) following Aurora B inhibition.

**Video 3: Fluorescent time-lapse imaging of telomere dynamics (Taz1-gfp) during meiosis II.**

Movie of a wild type cell expressing Taz1-gfp (telomeres, green) and Cdc11-cfp (SPBs, red).

**Video 4: Fluorescent time-lapse imaging of telomere dynamics (Taz1-gfp) during meiosis II following Aurora B inhibition.**

Movie of a wild type cell expressing Taz1-gfp (telomeres, green) and Cdc11-cfp (SPBs, red) following Aurora B inhibition in meiosis II.

**Video 5: Fluorescent time-lapse imaging of condensin dynamics (Cnd2-gfp) during meiosis I.**

Movie of a wild type cell expressing Cnd2-gfp (condensin, green) and Cdc11-cfp (SPBs, red).

**Video 6: Fluorescent time-lapse imaging of condensin dynamics (Cnd2-gfp) during meiosis I following Aurora B inhibition.**

Movie of a wild type cell expressing Cnd2-gfp (condensin, green) and Cdc11-cfp (SPBs, red) following Aurora B inhibition in meiosis I.

**Video 7: Fluorescent time-lapse imaging of condensin dynamics (Cnd2-gfp) during meiosis II.**

Movie of a wild type cell expressing Cnd2-gfp (condensin, green) and Cdc11-cfp (SPBs, red) in meiosis II.

**Video 8: Fluorescent time-lapse imaging of condensin dynamics (Cnd2-gfp) during meiosis II following Aurora B inhibition.**

Movie of a wild type cell expressing Cnd2-gfp (condensin, green) and Cdc11-cfp (SPBs, red) following Aurora B inhibition in meiosis II.

**Supplemental Table 1.**
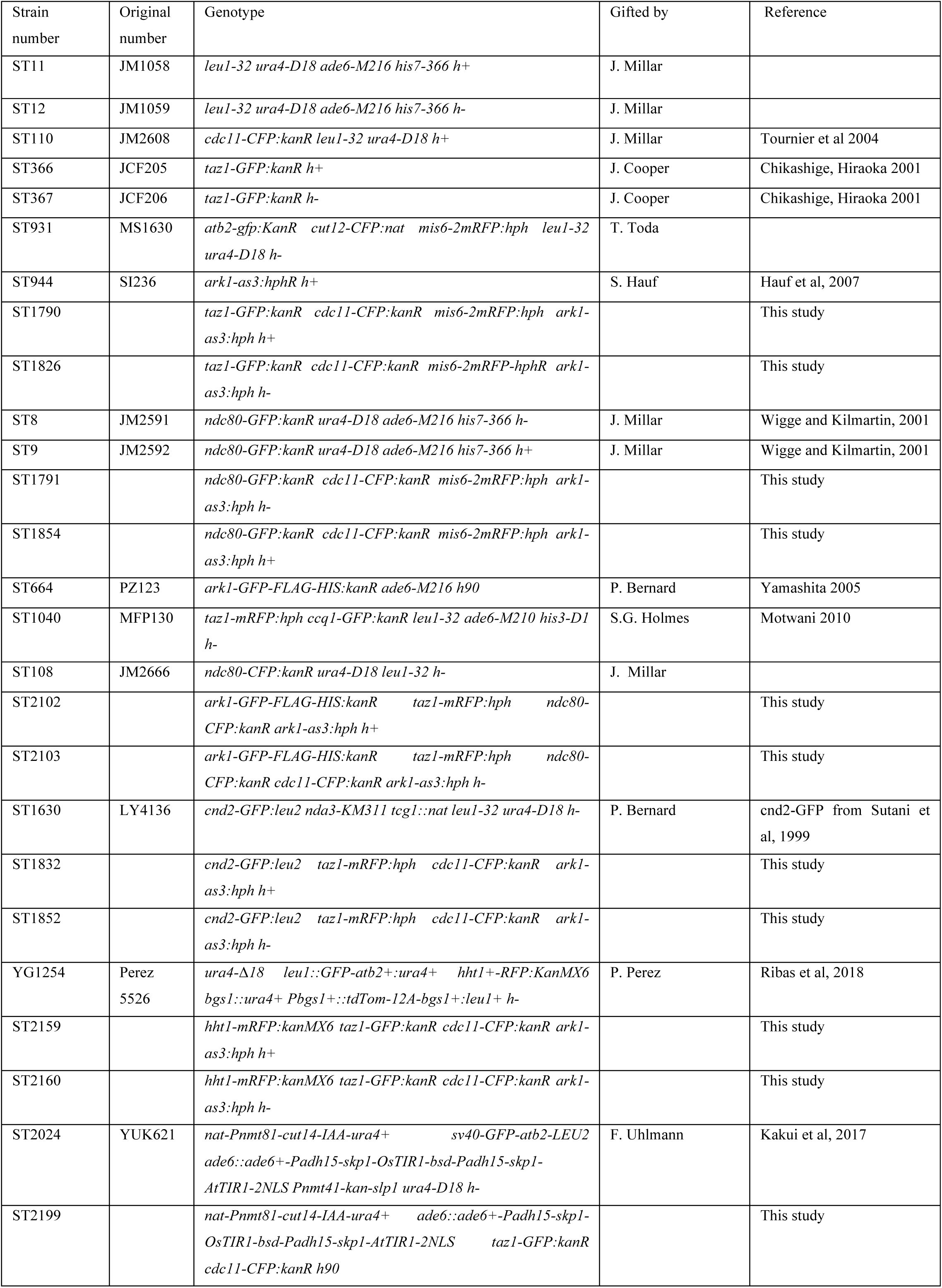
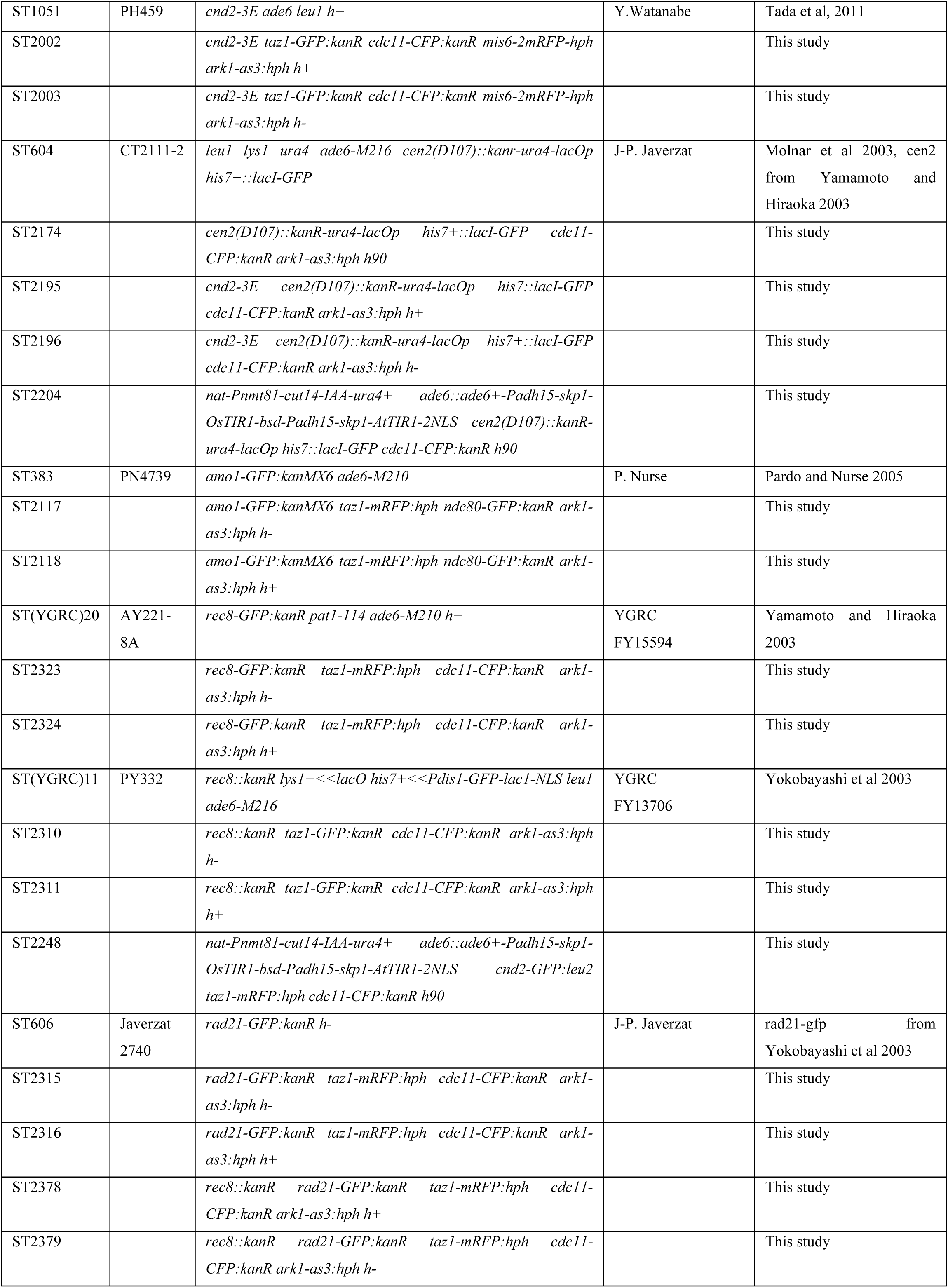
Strains used in this study.

